# Genome-scale metabolic modelling of trimethoprim’s mode of action reveals free ATP salvaged to restore purine pool

**DOI:** 10.1101/2025.07.25.666800

**Authors:** Cailean Carter, Dipali Singh, John Wain, Gemma C Langridge

## Abstract

Trimethoprim is a clinically important antibiotic for treating urinary tract infections yet its mechanism of killing remains elusive due to a cascade of effected metabolic reactions. Metabolites available in growth media can selectively counteract trimethoprim which affects the interpretation of results. We sought to understand the full scope of trimethoprim’s impact on *Escherichia coli* metabolism and how metabolite availability affects trimethoprim outcomes. We applied flux balance analysis on a genome-scale metabolic model of *E. coli* to simulate trimethoprim activity under bacteriostatic and bactericidal conditions. Our results suggested that in the absence of environmental purines or nucleosides, trimethoprim induces salvage of ATP to repair DNA. We experimentally validated the result with ATP bioluminescence screening of 96 clinical *E. coli* isolates. Therefore, the choice of growth media composition significantly changes the outcome of trimethoprim challenge and should be taken into consideration when interpreting trimethoprim’s activity.

**Graphical abstract:** 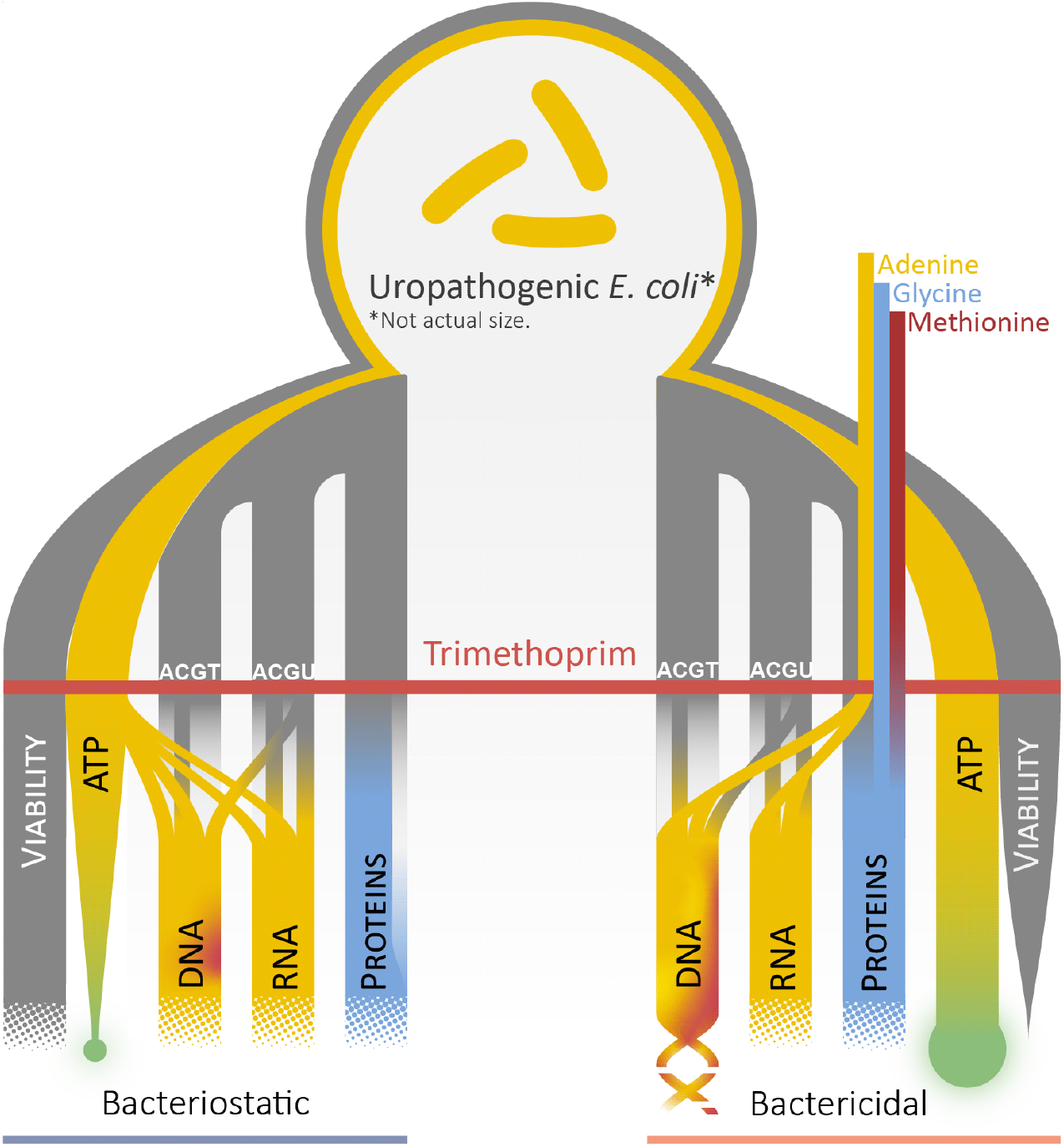

**Diagrammatic representation of trimethoprim action on susceptible UPEC in bacteriostatic and bactericidal conditions**. The diagram flows downwards in a time-irrespective manner with outcomes on cellular functions/properties not in order of events. The half-tone at the end represents slowing down of cellular process, however, the outcome is not clear in bactericidal conditions. Circular, green, glowing element at the end of ATP is a representation of ATP bioluminescence. Further, thinning of the line for viability and ATP denotes a reduction in the respective property. RNA and DNA components are labelled above the trimethoprim line with solid grey-to-yellow lines denoting components unaffected by trimethoprim activity. The components that are affected by trimethoprim are replenished by ATP, adenine, or uracil (U) and are denoted by the flow of lines. Abbreviations: A, adenine; C, cytosine; G, guanine; T, thymidine; U, uracil.

Sources: Kwon et al. ^1^, Amyes and Smith ^2^, Makino and Munakata ^6^, this study.

## Introduction

Urinary tract infections (UTIs) place a substantial burden on health care facilities and the quality of life for patients experiencing severe pain and discomfort ^3^. One antibiotic to treat UTIs, trimethoprim, is of clinical significance as the most cost-effective treatment option and is still used routinely despite rising resistance rates ^4,5^. Trimethoprim targets an important group of carbon-shuttling compounds called folates that are essential for providing carbon units for the synthesis of three of the four components of DNA (adenosine, guanosine, and thymidine), two of four components of RNA, and methionine (Figure 1) ^7^. Folates also provide the formyl group for protein synthesis initiatory *N-*formylmethionine (tRNA^fMet^). Therefore, trimethoprim directly affects DNA, RNA, and protein synthesis through limiting the availability of carbon-carrying folates.

**Figure 1.**
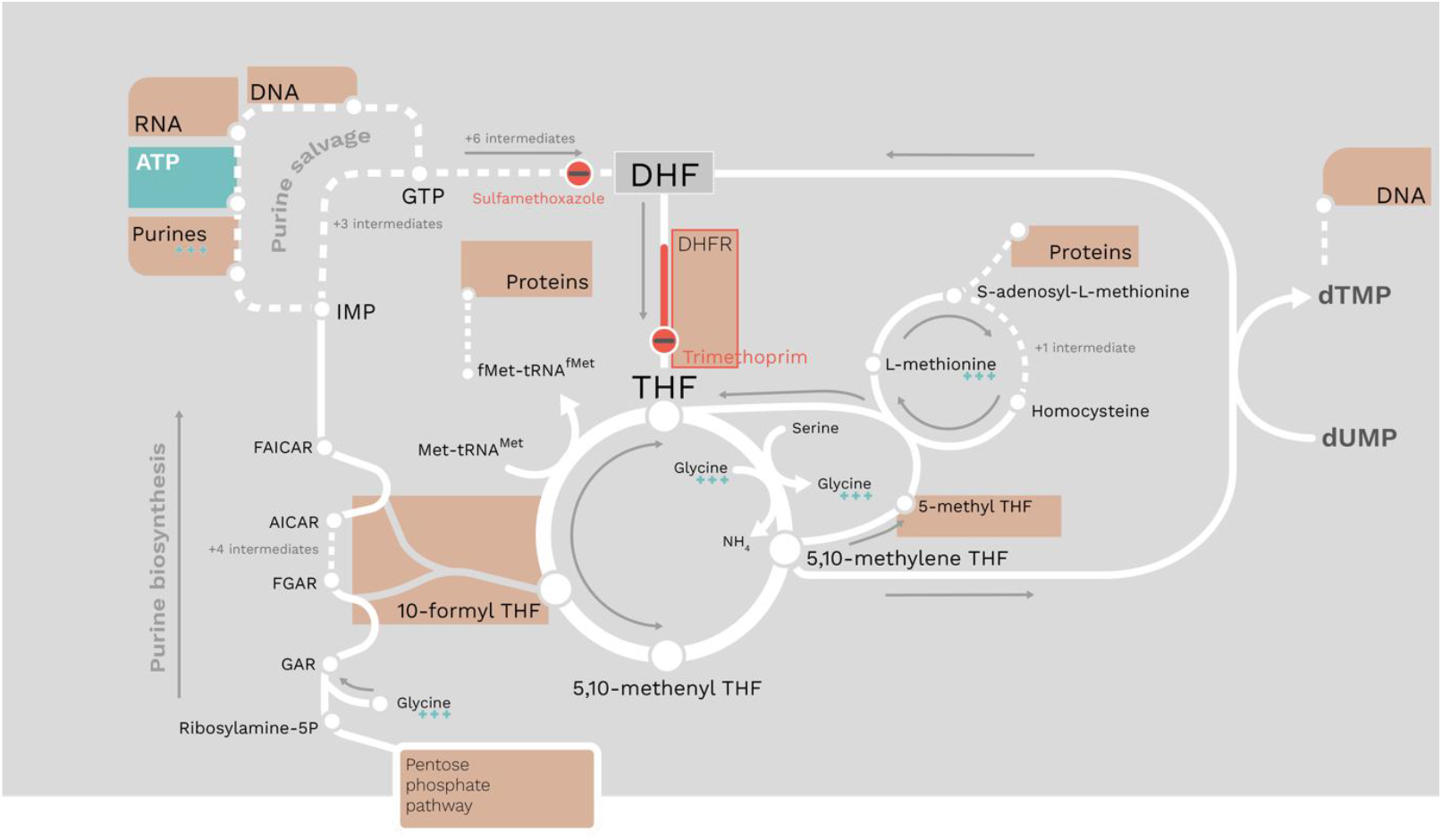
A simplified network map of the folate pool and relevant pathways including purine biosynthesis and salvage, and folate biosynthesis. White lines denote reactions and arrows for direction, sets of reactions have adjacent dark-grey arrows for direction. Dashed white lines denote several reactions and are condensed for clarity with labels stating number of intermediary reactions. Red circles with a black bar denote reactions where trimethoprim and sulfamethoxazole exert their anti-folate effects and metabolites/macromolecules/reactions with a light brown box denote components of metabolism affected by anti-folate drugs. Metabolites that are present in growth media which elicit a bactericidal activity are annotated with three teal plus signs. ATP is highlighted with the same teal colour to emphasise its position in the network. Dihydrofolate reductase (DHFR) is the only reaction labelled in the network map due to its significance as the target for trimethoprim. Reactions and pathways are derived from EcoCyc v21.

As these components of DNA, RNA, and proteins are also precursors for other metabolites, it is challenging to draw strong conclusions on the full extent of trimethoprim’s activity ^1^. This is further complicated by trimethoprim having bacteriostatic or bactericidal effects depending on metabolites available in growth media ^2^. Bacteriostatic trimethoprim activity depends on purines or nucleotides being excluded from growth media for trimethoprim to visibly slow or arrest cellular growth ^8,9^. Conversely, bactericidal activity requires the presence of a purine or nucleoside, and the amino acids glycine and methionine. The killing mechanism is suspected to be caused by inhibition of DNA synthesis via thymidine starvation while protein and RNA synthesis is alleviated. Faltered DNA synthesis and repair triggers programmed cell death via the suicide module *mazEF* and is analogous to thymineless death ^1,2,10,11^. The role of glycine is unknown with some suggestions that it plays a regulatory role in the bactericidal action but lacks sufficient evidence to rule out other amino acids that can replace glycine ^1,2^. Methionine replenishes trimethoprim-induced methionine depletion, thereby enabling protein synthesis which is essential for bactericidal activity ^2^. Therefore, a bactericidal outcome appears to require DNA, RNA, and protein synthesis via the availability of a purine, glycine, and methionine ^2,8^. Thymidine also plays an important role as a putative trimethoprim inhibitor; most organisms have a thymidine kinase to synthesise deoxythymidine monophosphate (dTMP) from thymidine which limits trimethoprim’s inhibition on DNA synthesis ^12^. The counteractive property of thymidine is only evident in bactericidal conditions yet metabolomics and protein expression analysis in bactericidal conditions suggests thymidine can only partially counteract trimethoprim ^1,2,13-16^.

As the full scope of trimethoprim’s activity on bacterial metabolism is unknown, we sought to better understand trimethoprim’s mode of action through genome-scale metabolic (GSM) modelling. The challenge with investigating folates and the trio adenine, glycine, and methionine is that they are highly connected metabolites. For example: adenine is involved with 20 reactions (excluding transporters), glycine with 49, and L-methionine with 56 (according to EcoCyc v27.1 *E. coli* K-12 substr. MG1655 database when accessed on 11/08/2023). Nonetheless, this challenge can be approached *in silico* with GSM modelling which provides a tractable representation of an organism’s metabolic network. Here, we focus on Uropathogenic *Escherichia coli* (UPEC), the predominant etiological agent of reported UTIs internationally; causing 77% of UTIs ^17^.

## Methods

### Strains and conditions

One hundred and ninety-nine clinical *E. coli* isolates from urine have previously been characterised with a further 18 collected for this study in 2020 using the same methods were collected from Norfolk & Norwich University Hospital ^18^. M9 minimal media (Formedium; MMS0101) was supplemented with 65 µg/mL adenine hemisulphate salt (Sigma), 50 µg/mL glycine (Fisher Scientific; G/0800/60), and 50 µg/mL L-methionine (Sigma; M-9625) as required. For all assays, isolates were grown in LB broth overnight at 37°C.

### Data analysis and visualisation

All data analysis was performed using the Python programming language (v3.8.10) with data handling on Jupyter Notebooks with the pandas (v1.1.3) and NumPy (v1.18.3) packages. All code is available on GitHub (github.com/CaileanCarter/Metabolic-modelling-trimethoprim-action). Euclidean and Cosine distances were performed using their respective method from the NumPy spatial distances module. All plots were drawn with ggplot2 (v3.4.2) using the R programming language (v4.2.1) with significance values plotted using ggsigniff (v0.6.4) and colour scheme provided by ggsci (v3.0.0) ^19^. Radar charts were plotted with the fmsb package (v0.7.5).

The metabolic network maps were created with the Escher Python package (v1.7.3) ^20^. The ScrumPy *E. coli* UTI89 model SBML file was converted into JSON format and all reaction and metabolite ids that were assigned a numerical value were changed to match the object’s name. The network map was drawn for the purpose of this study to encompass glycolysis, TCA cycle, pentose phosphate pathway, purine biosynthesis, purine salvage, one carbon pool by folate, electron transport chain, oxidative phosphorylation, and maintenance ATP import/export. It should be noted that reversibility data were lost in converting formats and all reactions are shown as reversible.

### Genome-scale metabolic modelling

Genome-scale metabolic (GSM) modelling was performed using ScrumPy3 (v3.0-alpha) and run on an Ubuntu 20.04 LTS on Windows Subsystem for Linux 2. An existing GSM model of *E. coli* UTI89 was used and is available on BioModels (MODEL2507240001). Model properties are described in the supplementary materials (Model properties and validation).

The media used for this model was M9 minimal media with individual transporters added for glucose, ammonium, sulphate, inorganic phosphate, water, oxygen, chlorine, and sodium. A list of biomass components produced by *E. coli* was derived from Feist et al. ^21^ which lists these components in proportion to each other. An exporting transporter was added for each biomass component. And the proportions were multiplied by the growth rate to obtain the flux values for biomass transporters (Table S1). For incorporating the metabolites supplemented to M9, constraints were added (when required) on the import of adenine, glycine, and methionine depending on the media condition being investigated. Where appropriate, flux bounds were applied to the import of these metabolites which represented the mM units of each that was supplemented to M9 minimal media. The media dependent constraints are outlined in Table 1.

**Table 1.**
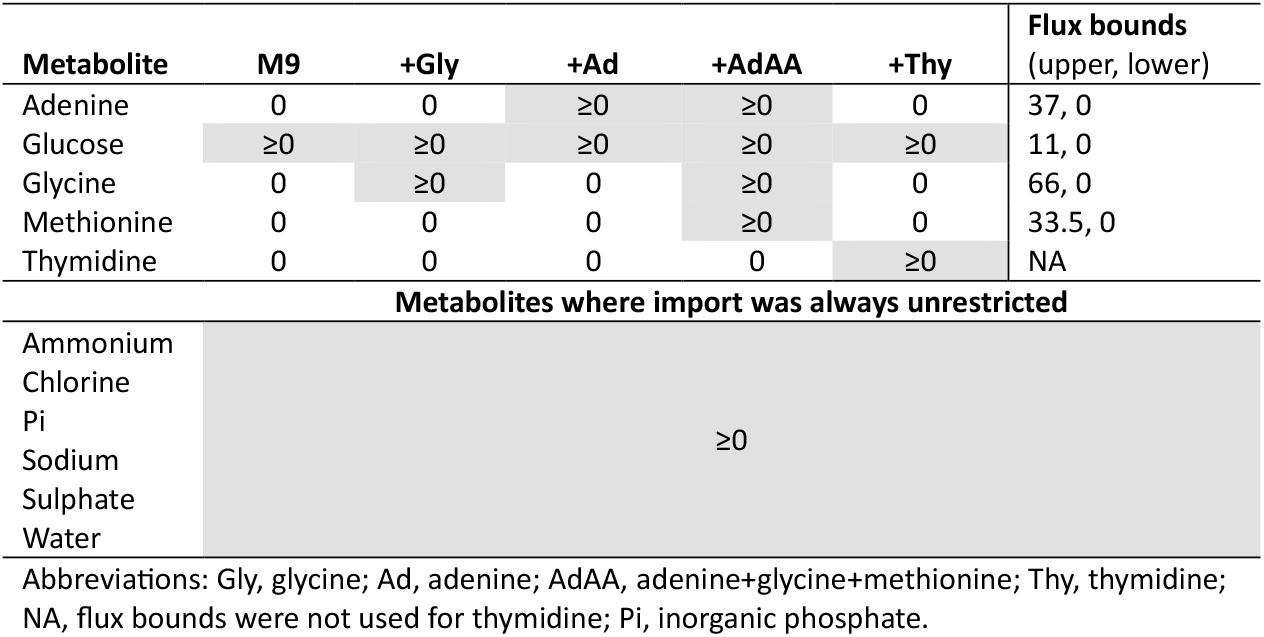
Constraints applied to import flux on media transporters for each media condition i.e., M9 minimal media with/without metabolite supplementation. For analyses that required flux bounds, the upper and lower flux bounds on import flux of that metabolite were used. Highlighted cells indicate metabolites available under each media condition.

### Folate associated reactions

A set of reactions concerning the folate pool was defined by identifying any reactions involving folate metabolites DHF, THF, 5-methyl THF, 5,10-methylene THF, 5,10-methenyl THF, or 10-formyl THF from the stoichiometry matrix of the *E. coli* UTI89 GSM (Table S5). The alternative formylated state of folate, 5-formyl-THF, was excluded as it is considered a storage form of folate and is not known to be a co-factor in any reaction (according to EcoCyc v27.1; accessed 03/03/2023). The biomass transporter for 5-methyl THF (MeTHF_bm_tx) was also excluded.

### Defining metabolic pathways

To ensure modularity and simplicity, the reactions most relevant to this project such as reaction involved in folate metabolism, electron transport chain (ETC), tricarboxylic acid (TCA), and purine metabolism were categorised into five distinct groups, as detailed in Table S4. The metabolic network map for these reactions is illustrated in Figure S3.

### Flux balance analysis

The methodology used here was as described previously ^22^. Briefly, flux balance analysis (FBA) was used for all metabolic modelling with minimisation of absolute total flux as the objective and steady state constraint (***N*** · ***v*** = 0). ***N***was a stoichiometry matrix of the *E. coli* UTI89 GSM with ***m*** rows for metabolites, ***n*** columns for reactions, and contents were the stoichiometries for each reaction. Further, ***v*** represented a vector of unknown flux values. Likewise, a weighting (ω) factor was sometimes changed for select reactions from the default value of 1 to confer a greater enzyme cost. Therefore, minimisation of the equation 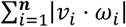.

### Identifying folate-dependency for biomass components

The production of each biomass component (Table S1) in the GSM was confirmed with a feasible linear programming (LP) solution producing one relative unit of said metabolite (Equation 1) and reported as absolute total flux of the solution (which was also the objective value).

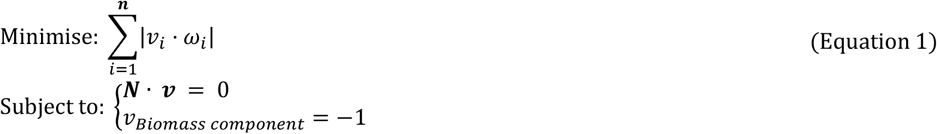

Absolute total flux of folates was calculated by summation of flux through folate associated reactions (Table S5) and provided a positive assertion that the synthesis of a metabolite utilised folates. The dependency on folates was determined using Equation 1 with the added constraint of setting flux for all folate associated reactions (Table S5) to zero while producing one unit of biomass component. The biomass component was deemed folate dependent if no feasible solution was made. To identify the reaction(s) within the folate pool used for the metabolite’s synthesis, a constraint of zero flux was added to Equation 1 for the synthesis of THF (DHFR), methionine (methionine synthase), purines (phosphoribosylglycinamide formyltransferase and AICAR transformylase), or 2-ketovaline (3-methyl-2-oxobutanoate hydroxymethyltransferase).

### Counteraction of trimethoprim-induced constraint on biomass synthesis by media supplementation

Counteraction by media supplementation was performed by permitting the import of adenine, glycine, methionine, and/or thymidine (Table 1) while all folate associated reactions (Table S5) were set to flux of zero as shown below:

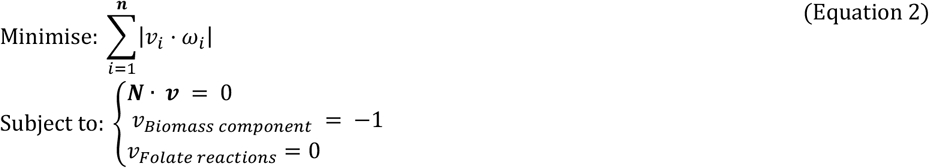

### Network response to increasing dGTP demand under trimethoprim-induced purine biosynthesis inhibition

To investigate how metabolite supplementation could alleviate trimethoprim-induced purine biosynthesis inhibition, a series of LP formulations were solved repeatedly with increasing flux for dGTP export for each media condition. Uniquely to this analysis, adenine alone was included as a supplementation condition. The LP formulation was made as described in Equation 3 and relevant media condition flux bounds constraints (Table 1) were applied (including glucose flux bounds). Arbitrary upper flux constraints were applied to two purine biosynthesis reactions involving folates, GAR formyltransferase and AICAR transformylase, to simulate trimethoprim restriction on purine biosynthesis. Export flux of dGTP was incremented to the maximum arbitrary value of 20; this upper bound needed to be greater than the bounds placed on purine biosynthesis but not so large that most of the output had no change in reaction sets. Limitations were placed on ATP salvage by restricting ATPase to a flux of zero.

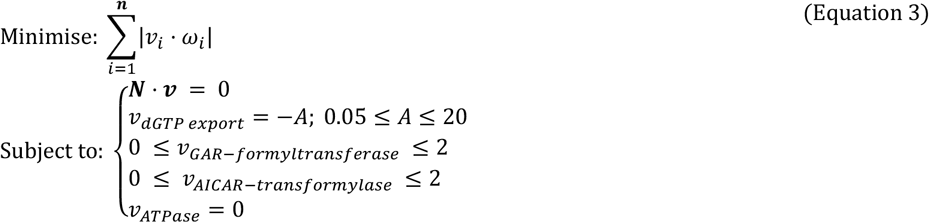

### ATP bioluminescence time-course assay

250 µL of overnight culture of a trimethoprim susceptible UPEC isolate was transferred to 25 mL pre-warmed media in a 250 mL baffled flask and incubated at 37°C, shaking at 140 rpm till an OD_600_ of ∼0.3. When solely using M9 without supplementation, 1 mL of overnight was transferred to 24 mL M9 to reduce the time taken to reach OD_600_ of ∼0.3. From the baffled flask, cells were diluted to 0.02 OD_600_ in 8 mL pre-warmed M9 media (supplemented when required) and vortexed thoroughly. 80 µL of diluted culture was transferred to each well of a 96-well Thermo Scientific Nunc optic clear bottom white walled plate. The assay was performed on the CLARIOstar Plus (BMG Labtech) with two injectors using a custom script provided by BMG Labtech which was further modified to meet the experimental design described here. One injector was primed with BacTitre-Glo reagent (Promega) and prepared according to manufacturer’s specifications. The CLARIOstar Plus’ environmental control was set to 37°C for the duration of the assay. Trimethoprim (Sigma-Aldrich; SBR0024) was prepared at 9X the desired concentration in pre-warmed M9 media (supplemented when required). The assay began with a 5-minute incubation period before measuring OD_600_ across the whole plate every 15 minutes for three measurements. Each column of the plate was treated as a timepoint in the assay. Following the three baseline absorbance measurements, three ATP bioluminescence baseline measurements were taken. For each timepoint, the OD_600_ was measured first before injecting an equal volume in the well of BacTitre-Glo reagent and shaking at 100 rpm for a minute, and measuring ATP bioluminescence using the Ultra-Glo pre-set. The plate was then ejected and 10 µL of pre-warmed media (control) or antibiotic (treated) solution was added to the appropriate wells and inserted back into the machine. The remainder of 15 minutes was waited for. Thereon, ATP bioluminescence readings were taken every 10 minutes for nine measurements. All time-course assays were repeated in triplicate.

### Incorporating experimental data to model the salvage of free ATP under trimethoprim stress

To better understand how free ATP would be used under trimethoprim challenge, experimental data were incorporated into the model whereby the network would synthesise free ATP and a portion of this could later be salvaged under trimethoprim challenge. Although *in vivo* free ATP would be available intracellularly within the population of cells as the population grew, *in silico* FBA under steady state does not permit accumulation of intracellular metabolites (hence biomass components are treated as external metabolites). Therefore, an external ATP pool was introduced that could be salvaged by the network under conditions of stress. This form of ATP was termed maintenance ATP (mATP) to differentiate it from the biomass ATP which was a component of RNA which hereon is referred to as biomass ATP (Table S2).

### Converting optical density to cell counts for growth rate

The absorbance (*OD*) data were converted into total number of cells (*n*_*cells*_) for each value at each time point (*t*) using the formula described in Equation 4. Absorbance was converted to total cell counts for two reasons: a) there was a change in well volume during the assay (addition of media/trimethoprim) which increased the distance light needed to travel through the liquid which alters OD and b) the growth rate derived from absorbance did not scale biomass export into the same range as the ATP bioluminescence measurements. Total cell count was calculated with a conversion factor from Volkmer and Heinemann ^23^ for *E. coli* in M9 minimal media with glucose supplementation (Equation 4). This conversion factor was multiplied by the volume (*V*) of the well and OD.

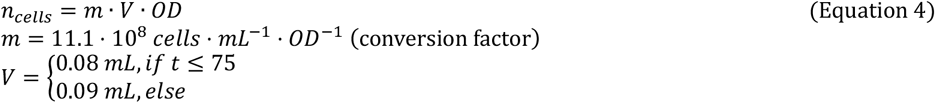

### Linear regression

The cell numbers and ATP bioluminescence data from the time-course assays were used to provide the growth rate and total ATP for the metabolic model for each media and treatment condition. A linear regression was made using the Python package statsmodels’ *ols* function (v0.12.2) and included baseline measurements from control groups. Baseline measurements for treated groups were excluded from the models as control groups would be used instead. Separate linear regressions were made for cell counts and ATP bioluminescence (FLU), and each included time, media type, and treatment as predictive variables with interactions between them (R formula: [*n*_*cells* |_ *FUL*∼*time* * *C*(*treatment*) * *C*(*media)*). Assumption were checked using the Python package Seaborn *lmplot* for linearity, a qqplot to assert data were normally distributed, and a stem plot of Cook’s distance to assert no data points had undue influence over the model. The growth rate (*µ*) was calculated for each media and treatment group from the linear regression; the slope of the line from the cell count model was multiplied by a constant factor of 0.1 from Díaz Calvo et al. ^24^ which enabled comparisons between OD and growth rate.

### Total free ATP, maintenance ATP, and biomass ATP

A maintenance cost normally covers the energy required to maintain basic cell functions like cell membrane integrity, polymerisation of proteins, and repairing DNA, however, the maintenance costs were unknown for the different media and treatment conditions. Further, it was suspected that this maintenance cost would be dramatically different between these different conditions ^25^. Therefore, experimental values were used to capture the total free ATP which included mATP. Although it was suspected that the demand for RNA and DNA components would change immediately after trimethoprim challenge, experimental data was not available. Thus, the demand for these components immediately after trimethoprim-like conditions were the same as control conditions, which was proportional to the growth rate.

Experimentally collected ATP bioluminescence data were used as an indicator of total free ATP present in the population for each treatment and media condition. As there was no available conversion factor for converting relative light units (RLUs) derived from ATP bioluminescence to *in silico* total ATP, it was assumed that RLUs equalled the total ATP present in a sample. This was informed by the reaction stoichiometry for the luciferase reaction (Table S6) used for ATP bioluminescence; the net stoichiometry implied that *in vivo* total free ATP is directly proportional to the number of photons, measured as RLUs, which was measured *in vivo*.

The total free ATP captured by the luciferase reaction was assumed to capture both mATP and biomass ATP (but not ATP already incorporated into RNA). Experimental RNA synthesis rates were not collected, therefore, the rate of ATP produced for RNA in each media and treatment condition was the flux value of RNA ATP export (Table S1) multiplied by the growth rate (*µ*). Likewise, the rate of change in total free 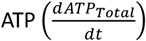 for each media and treatment condition was the slope of the linear regression line. Therefore, the unknown mATP cost was calculated by subtracting the biomass ATP production rate from the total free ATP production rate.

### Defining a salvageable external maintenance ATP pool

To observe whether the model would salvage mATP under trimethoprim stress in the different media conditions, a pool of external mATP was made available to the model. Like *in vivo*, the model needed to create this pool of mATP as the population grew under normal growth conditions before trimethoprim stress was evoked. Therefore, the quantity of mATP in a sample was proportional to the population size. However, as biomass ATP and mATP were unknown quantities, their rates of change were used to determine proportions as described in Equation 6. Therefore, the relative units (RU) of mATP change over time could act as a proxy for the availability of mATP for salvage. Also, flux values for total ATP were used rather than absolute values to see how mATP was produced and used under trimethoprim stress, rather than telling the model how much mATP needed to be salvaged.

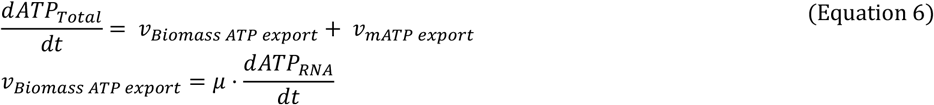

To assert there would be sufficient demand for ATP relative to the mATP available, a series of calculations were solved to find the proportions of biomass ATP and mATP at each time point from the ATP time-course assays (Equation 7). These calculations did not require FBA as internal flux values were not needed for this analysis. Because the assay measurements began at the 30-minute point, 1 minute was used as the time difference at the 30-minute mark. Baseline measurements for the treated group were from the control group as the linear regression excluded baseline measurements as it would not have passed the linearity assumption.

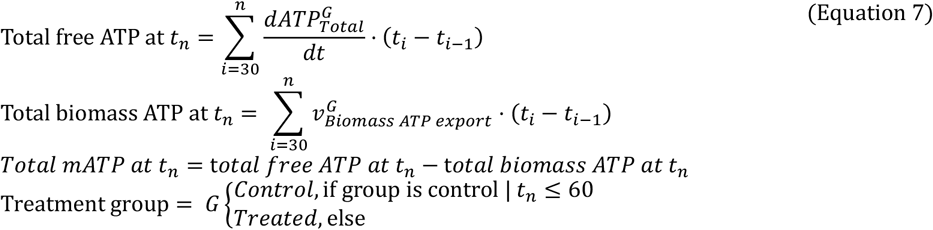

### Optimum weighting for mATP import and folate associated reactions

To be confident that the model would not import mATP as a free energy source, an optimum weighting was required for the mATP import reaction (mATP_tx) which would prevent the model importing mATP under normal growth conditions. A series of LP solutions with the base formulation in Equation 8 were calculated with increased weighting applied to mATP import. This was done for each media condition as media supplementation meant different energy requirements, therefore, different optimum weightings. The optimum weighting was determined to be the minimum mATP_tx weighting whereby flux through mATP_tx was zero.

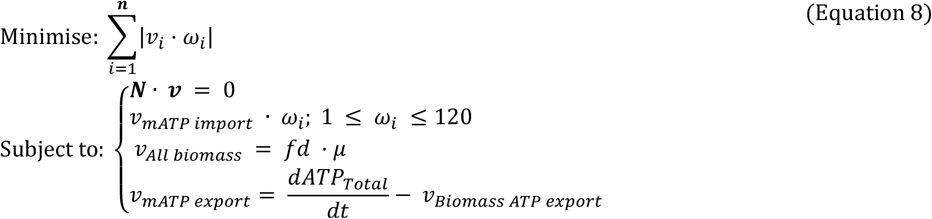

To simulate trimethoprim conditions, an optimum weighting was required to be applied to all folate associated reactions (Table S5), for all media conditions (Table 1), that would result in the import of mATP. This was performed similarly to mATP_tx and included media-specific optimum weighting for mATP import 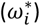 as determined in Equation 8. The optimum weighting for folates was deemed to be the minimum weighting on folate associated reactions which achieved the maximum import flux of mATP. A summary LP formulation is provided in Equation 9.

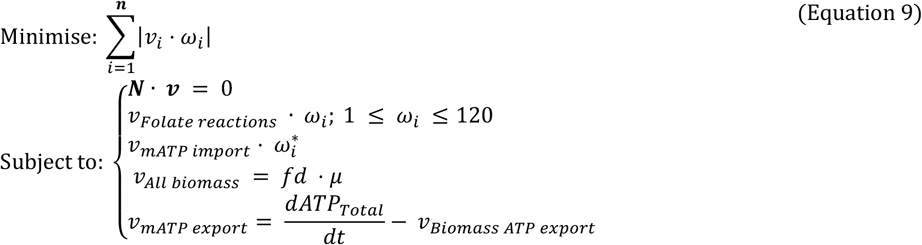

## In-silico time-course ATP assay

The *in-silico* ATP time-course assays were simulated by enabling the model to salvage external mATP. Because the LP formulation for this analysis required some mATP to be available at the start, omics data were used instead of an arbitrary number as it was not possible to do a zero division. Thus, prior to the first time point (<30 minutes) the availability of mATP was calculated using Equation 7 with the addition of the intercept from the ATP bioluminescence linear regression for each condition. For example, to calculate the total free ATP before baseline measurements:

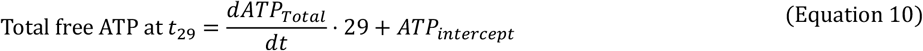

From thereon, a series of LP formulations were solved at each time point (*t*). The import of mATP was constrained to the total accumulated mATP divided by Δ*t* to prevent the solution from importing more mATP than was available in the external mATP pool. Constraints applied during baseline time points for the treated group were the same as the control group which included growth rate and weighting applied to folate associated reactions. At the 60-minute mark, the weighting to folate associated reactions was applied to the treated group, however, the control group’s growth rate was used until the 75-minute mark to simulate the demand for ATP. This is summarised in the following LP formulation:

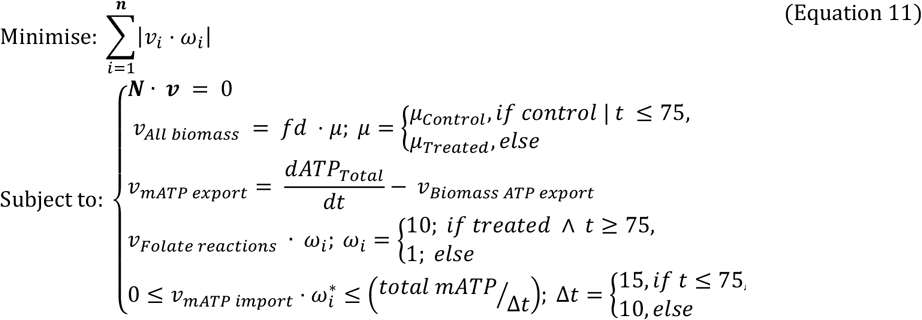

The total external ATP was calculated at each time point using Equation 12 with the division of biomass ATP and maintenance ATP following the principle in Equation 6.

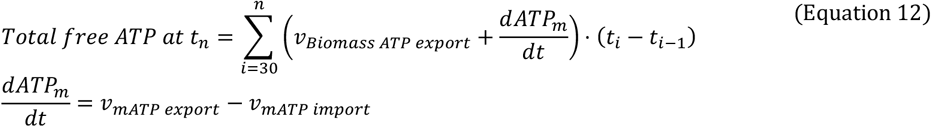

The centrality of reactions of reactions was calculated using PageRank (*PR*) with a dampening factor of 0.85 ^26^.

### Mass flow graphs and functional groups

For clustering reactions into functional groups to understand glycine’s role in bactericidal trimethoprim action, mass flow graphs were made using a modified approach from Beguerisse-Díaz et al. ^27^. For each media and treatment group permutation, the stoichiometry matrix (***N***) was filtered for reactions and metabolites present in the LP solution. The consumption and production of each metabolite per reaction in ***N***was calculated by multiplying the respective flux value (***v***) from the LP solution by the column (***n***), giving ***N***(***v***). An adjacency matrix (***A***_***ij***_) was made from ***N***(***v***). For calculating the edges, the flow of a metabolite 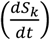 from a producing reaction (*R*) to a consuming reaction (*R*_*j*_) was calculated using an adjusted equation from Beguerisse-Díaz *et al*. ^27^ as shown below:

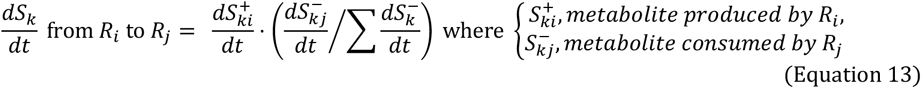

The summation of Equation 13 across all metabolites connected between *R*_*i*_ and *R*_*j*_ were used as weighting (***ω***) for the edges in ***A***_***ij***_:

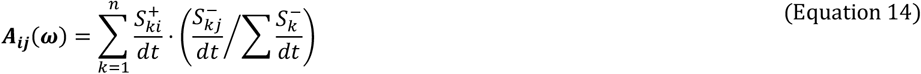

Mass flow graphs were drawn with igraph (v0.10.6) in Python using the edges determined above to draw connections between metabolites with mass flow as a weighted attribute. The shared community structure was identified using the Leiden algorithm Python package (v0.10.1) with the *find_partition_multiplex* method and arguments ModularityVertexPartition, number of iterations of - 1 for the algorithm to run until no improvement was made, seed of 1, and weights were set to the edge weights ^28^.

### High-throughput ATP bioluminescence screening

Overnight cultures were performed in clear 96-well plates with 300 μL LB broth which were inoculated from frozen stocks. 8 μL of overnight culture were transferred to two wells with 72 μL of M9 media (supplemented when required) in white 96-well plates (Thermo Scientific). Cells were left to acclimate for 10 minutes at 37°C. 10 μL of M9 media (supplemented when required) was added to control wells and 10 μL 36 mg/L trimethoprim was added to treated wells for a final concentration of 4 mg/L and incubated at 37°C for 30 minutes. ATP bioluminescence was measured using the CLARIOstar Plus. For this, the plate reader injected 90 μL of BacTitre-Glo to each well followed by shaking at 100 rpm for 20 seconds. Fluorescence was measured at 550-570nm one minute after injection. The change in fluorescence between untreated and treated wells was calculated by *RLU*_*treated*_*/RLU*_*untreated*_ and taking the mean across triplicates.

### Statistical analysis for ATP bioluminescence screening

A binomial logistic regression was used to determine whether the change in fluorescence between treated and untreated in trimethoprim susceptible and resistant samples were statistically significant in each media condition. This was performed using the *glm* method from *statsmodels* (using binomial family) with change in fluorescence as the predictor variable and trimethoprim susceptibility as the endogenous variable. The linearity between the predictor variable and logit was checked before calculating the log odds. A stem plot of Cook’s distance was also used to assert no data points had undue influence over the model. Receiver operating characteristics (ROC) were performed by iterating over a series of threshold values between 1% and 300%. Optimal cut-off values for distinguishing susceptibility were identified by maximum Youden’s index.

### Variability in ATP between ATP bioluminescence screening

For comparing deviation from the sampling mean, the change in fluorescence for each replicate of an isolate was adjusted by subtracting the isolate mean: 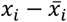 whereby *x*_*i*_ was the change in fluorescence for each replicate measurement for each isolate, 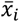 was the mean change in fluorescence for the isolate *x*_*i*_. Therefore, the overall population mean was zero – allowing straightforward comparison in deviation from the mean between isolates. Two standard deviations (±2σ) of all adjusted data points within a group were then calculated.

## Results

### Trimethoprim disruption of biomass synthesis in bacteriostatic conditions

Trimethoprim is known to inhibit the synthesis of some amino acids and purines; therefore, it was prudent to replicate this *in silico* with a genome-scale metabolic (GSM) model (Equation 1). By mimicking trimethoprim’s constraints on folate reactions *in silico*, we found trimethoprim directly affected the synthesis of 15 biomass components (excluding derivates like NAD:NADH): 8 purine and (deoxy)nucleotide triphosphates, 2 amino acids, 3 cofactors, one amine compound, and folates (Figure 2a).

**Figure 2.**
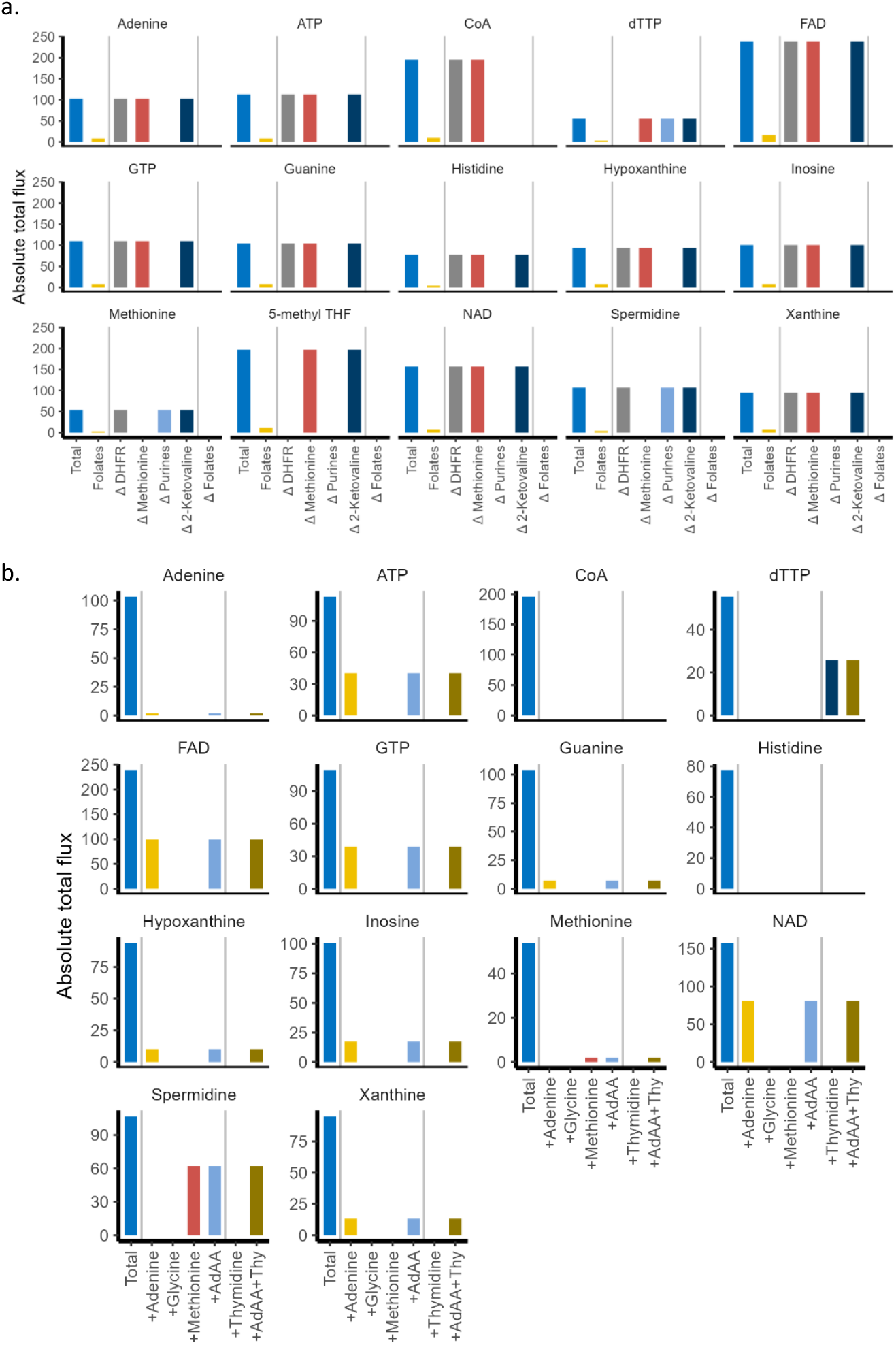
Biomass components affected by trimethoprim-induced disruption of the folate pool and counteraction by adenine, glycine, methionine and/or thymidine. (A) Total denotes the absolute total flux through the network to produce 1 unit of metabolite. The Folates bar (yellow) indicates the proportion of which involved folate associated reactions (Table S5). Labels on x-axis with Δ denote the total flux through the network when a reaction in the folate pool was restricted to zero flux in the presence of a given metabolite. This includes: dihydrofolate reductase (ΔDHFR); methionine synthase (ΔMethionine); GAR formyltransferase and AICAR transformylase (ΔPurines); dehydropantoate hydroxymethyltransferase (Δ2-Ketovaline). (B) Impact of adenine, glycine, methionine, and/or thymidine supplementation on metabolites which required folates for their synthesis when folate associated reactions are constrained. The total denotes the absolute total flux to synthesise the given metabolite when no constraint was applied to folate associated reactions and without supplementation. For each metabolite being supplemented, the flux for all folate associated reactions was set to zero and shows the absolute total flux of the LP solution. Abbreviations: adenine, glycine, and methionine (AdAA); thymidine (Thy).

This analysis highlighted some acute differences between what would be expected *in vivo* and how to reproduce it *in silico*. Most notably, preventing flux via DHFR (the target enzyme for trimethoprim) only affected the synthesis of folates (in the form of 5-methyl THF) and dTTP. It was then suspected that some biomass components utilised folates but do not require DHFR for their synthesis under steady state. Indeed, the other biomass components that required folates were dependent on the folate ‘pool’ rather than DHFR activity. This implied that the folate pool had a conserved moiety (a group of compounds that be cyclically interconverted to one another) as THF could acquire a carbon unit from serine or glycine, donate that to the synthesis of a metabolite, and revert to THF. This also made investigating trimethoprim a challenge as FBA does not allow for the depletion or accumulation of internal metabolites under steady state. A second concern was that folates could not be salvaged in the model. Therefore, when synthesising folates *de novo*, the only means of balancing the solution was by exporting folates as 5-methyl-THF which is one of the biomass components (Table S1).

Nonetheless, the biomass components that directly required folates for their synthesis were identified. Of particular interest was ATP which required folates for *de novo* synthesis via enzymes involved in the purine biosynthesis pathway that utilised 10-formyl THF. The dependence on folate in histidine and spermidine synthesis was unexpected as this is not well reported in the literature. However, histidine was derived from parts of the purine biosynthesis pathway that involved folates (see review by Winkler and Ramos-Montañez ^29^). Upon closer inspection, the biosynthesis of spermidine relied on S-adenosylmethionine (SAM) which required folates for its biosynthesis via synthesis of L-methionine ^30^. All purines (and their nucleoside [triphosphate] counterparts) and cofactors NAD, FAD, and CoA required folates for their biosynthesis via GAR formyltransferase and AICAR transformylase (Figure 2a).

### Supplementation with adenine, glycine, methionine, or thymidine counteracts trimethoprim selectively

To better understand the relevance of adenine, glycine, and methionine in eliciting bactericidal trimethoprim activity, and how thymidine counteracts trimethoprim, their capacity to counteract trimethoprim-induced constraint on biomass synthesis was investigated (Equation 2). Under the condition of zero flux via folate-associated reactions (Table S5), adenine supplementation permitted the synthesis of all purines and nucleosides (Figure 2b). Cofactors FAD and NAD were also restored as GTP was the precursor for FAD as described in the “flavin biosynthesis I (bacteria and plants)” pathway in EcoCyc v27.1 and NAD required ATP for its synthesis. This suggested adenine restored the purine, ATP, and cofactor synthesis under trimethoprim challenge. In contrast, the synthesis of CoA was not restored by supplementation as the constraint applied to dehydropantoate hydroxymethyltransferase (one of the folate-associated reactions) restricted synthesis of 2-dehydropantoate which is a precursor for CoA. While histidine and purines share a common pathway for their synthesis, which require folates, the supplementation of adenine did not restore histidine synthesis. Glycine alone did not restore the synthesis of any biomass components (Figure 2b).

Further, the presence of glycine with adenine and methionine did not alter the absolute total flux for synthesising any biomass components compared to adenine or methionine alone. Thus, glycine’s relevance in the activity of trimethoprim or as a lone supplement was not immediately clear from this analysis. Methionine restored the synthesis of itself and spermidine for which methionine is a precursor (Figure 2b). No biomass components were exclusively restored by the combination of adenine, glycine, and methionine which suggests these metabolites selectively counteract trimethoprim. Thymidine allowed the synthesis of dTTP when all folate associated reactions were void (Figure 2b). Indeed, none of the bactericidal promoting metabolites altered the total flux required to synthesise dTTP when thymidine was present. Thus, thymidine alone counteracted the limitation applied to thymidylate synthase under trimethoprim action. As the reactions involved in synthesising folates were constrained to zero, the biomass component 5-methyl THF was excluded for this analysis.

### Purine supplementation restores the nucleoside pool during trimethoprim-induced constraints on purines biosynthesis

The capacity for a purine to restore the nucleoside pool under trimethoprim challenge was investigated by increasing nucleoside demand (dGTP export) with constraints applied to purine biosynthesis reactions involving folates (GAR formyltransferase and AICAR transformylase). To simulate trimethoprim’s activity on purine biosynthesis, an upper constraint was applied to these two reactions as a proxy for limited active folate availability (Equation 3). Therefore, the aim was to observe how the model responds when dGTP demand exceeds the constraints applied to purine biosynthesis i.e., whether the model can still synthesise dGTP or rely on media supplementation. In bacteriostatic conditions without purine availability from the media, the model used purine biosynthesis to synthesise dGTP *de novo* until the upper bounds placed on purine biosynthesis were met (Figure 3). Past those bounds, no feasible solution was available for synthesising purines. Glycine supplementation alone was not sufficient for surpassing the bounds, but adenine supplementation enabled the model to continue making dGTP through adenine salvage until the limited availability of glucose became restrictive. The same applied to bactericidal conditions, however, the availability of glycine as a carbon source enabled the model to reach double the maximum flux of dGTP than with adenine alone. Also, methionine was not imported in bactericidal conditions, suggesting methionine was not used as a carbon source in this condition.

**Figure 3.**
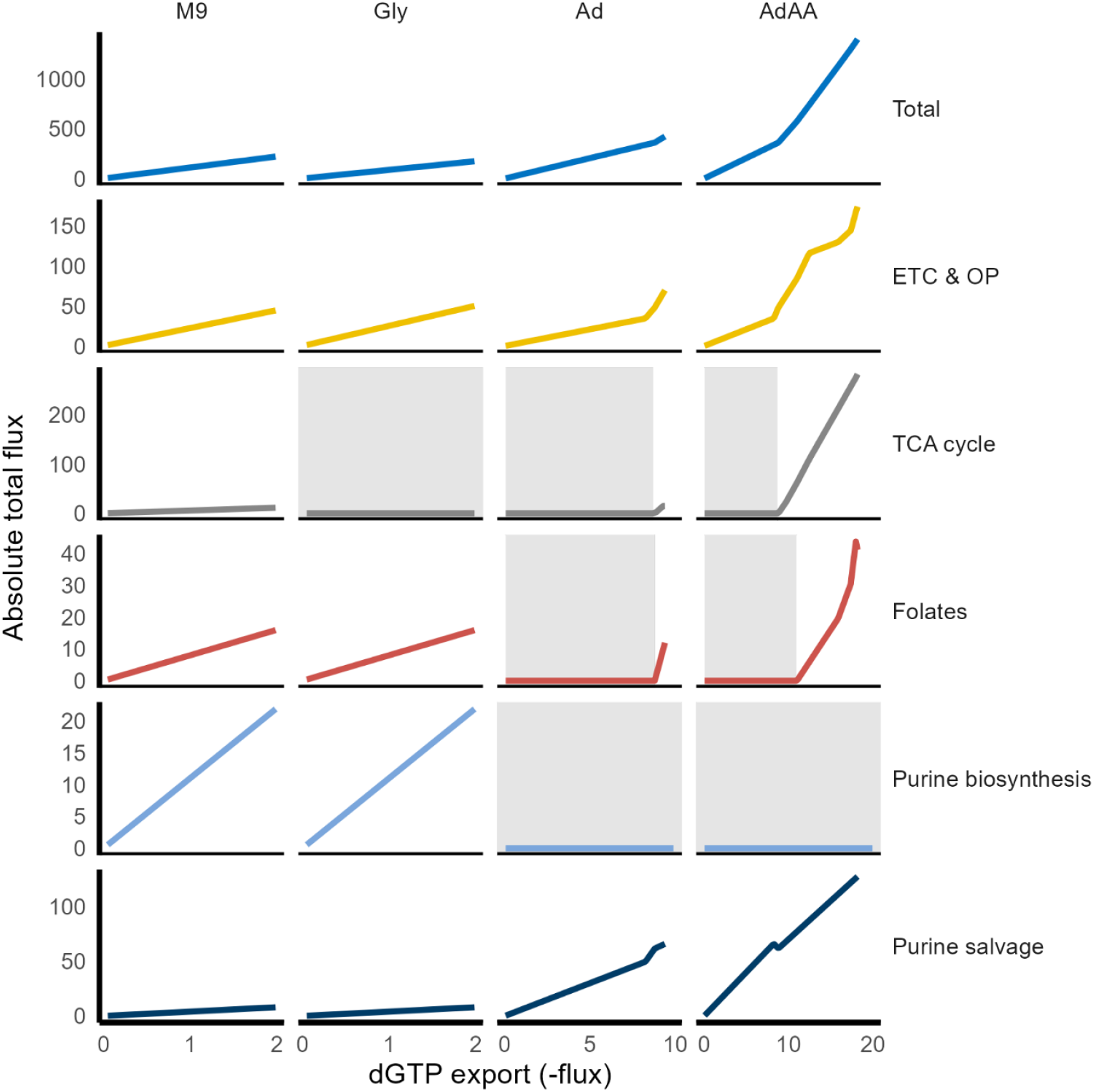
Network response to increasing dGTP demand under trimethoprim-induced *de novo* purine biosynthesis limitation in M9 minimal media with supplementations of glycine (gly), adenine (Ad) or adenine, glycine, and methionine (AdAA). Each line represents the absolute total flux through a pathway as the flux for dGTP export is increased. Purine salvage under AdAA conditions contains three lines representing the switching of routes as demand increased. Flux via GAR formyltransferase and AICAR transformylase was restricted to a maximum of 2 to imitate trimethoprim-induced purine biosynthesis limitation. Grey backgrounds denote dGTP fluxes where no flux was found for that pathway. Abbreviations: tricarboxylic acid (TCA) cycle; electron transport chain (ETC) and oxidative phosphorylation (OP).

### Incorporating *in vivo* ATP omics data suggests ATP is used to restore the nucleoside pool during bacteriostatic trimethoprim action

*In vivo* absorbance and fluorescence data were integrated into the model to assess how the network utilises free ATP, measured by bioluminescence and partitioned into growth-related (biomass ATP) and maintenance ATP (mATP) (Figure S4a&b). Akin to other biomass components, the biomass ATP was directly proportional to the growth rate. Assuming that mATP could only be salvaged under trimethoprim stress, we incorporated weightings for mATP import in different media compositions to prevent its use during normal growth. Trimethoprim’s activity was simulated by applying weightings to folate-associated reactions which (unlike directly inhibiting DHFR) allowed for feasible solutions (Equation 8 & 9, Figure S4c). Preliminary observation by Kwon *et al*. ^1^ reported that glycine supplementation into M9 minimal media induces a rapid decline in ATP immediately after trimethoprim exposure in *E. coli* and so we included glycine supplementation for this analysis.

We then simulated how the network salvaged mATP under trimethoprim stress in each media condition (Equation 11). In bacteriostatic conditions, mATP was imported after 60 minutes in trimethoprim-like conditions (Figure 4a). The simulated decline in total external ATP for treated groups compared to control at 75 minutes mirrored the proportion of ATP decline observed *in vivo* (− 32.9% in M9 and -35.9% in M9+Gly; Figure 4b) whereas there was no decline in total ATP in bactericidal conditions. Unlike M9 conditions where the treated group observed a single large shift in the proportion of mATP (at time of trimethoprim constraint) before a maintained mATP proportion, there was a constant decline in the %mATP in M9+Gly conditions. By 125 minutes, mATP accounted for 0.24% of total ATP produced in M9+Gly compared to 36.6% in M9 conditions. In bacteriostatic conditions, a significant portion of imported mATP flux could be attributed to the export flux of mATP (36% in M9 and 25.9% in M9+Gly). Of the flux that was not immediately exported as mATP in both bacteriostatic conditions, 45.8% could be attributed to the export flux of biomass ATP and dATP, and 52.9% attributed to export flux of GTP and dGTP. In bactericidal conditions, 12.4% of adenine import could be attributed to the export of mATP, and the remaining adenine was used to produce the same percentages of purines and nucleosides seen in bacteriostatic conditions. This strongly suggested that purines and purine-based nucleosides were the greatest contributor for import of mATP and adenine. Further, purine salvage was a more central component of the network in bacteriostatic media conditions in treated groups compared to the control group, suggesting the ATP salvaged was distributed across the network (Figure 4d).

**Figure 4.**
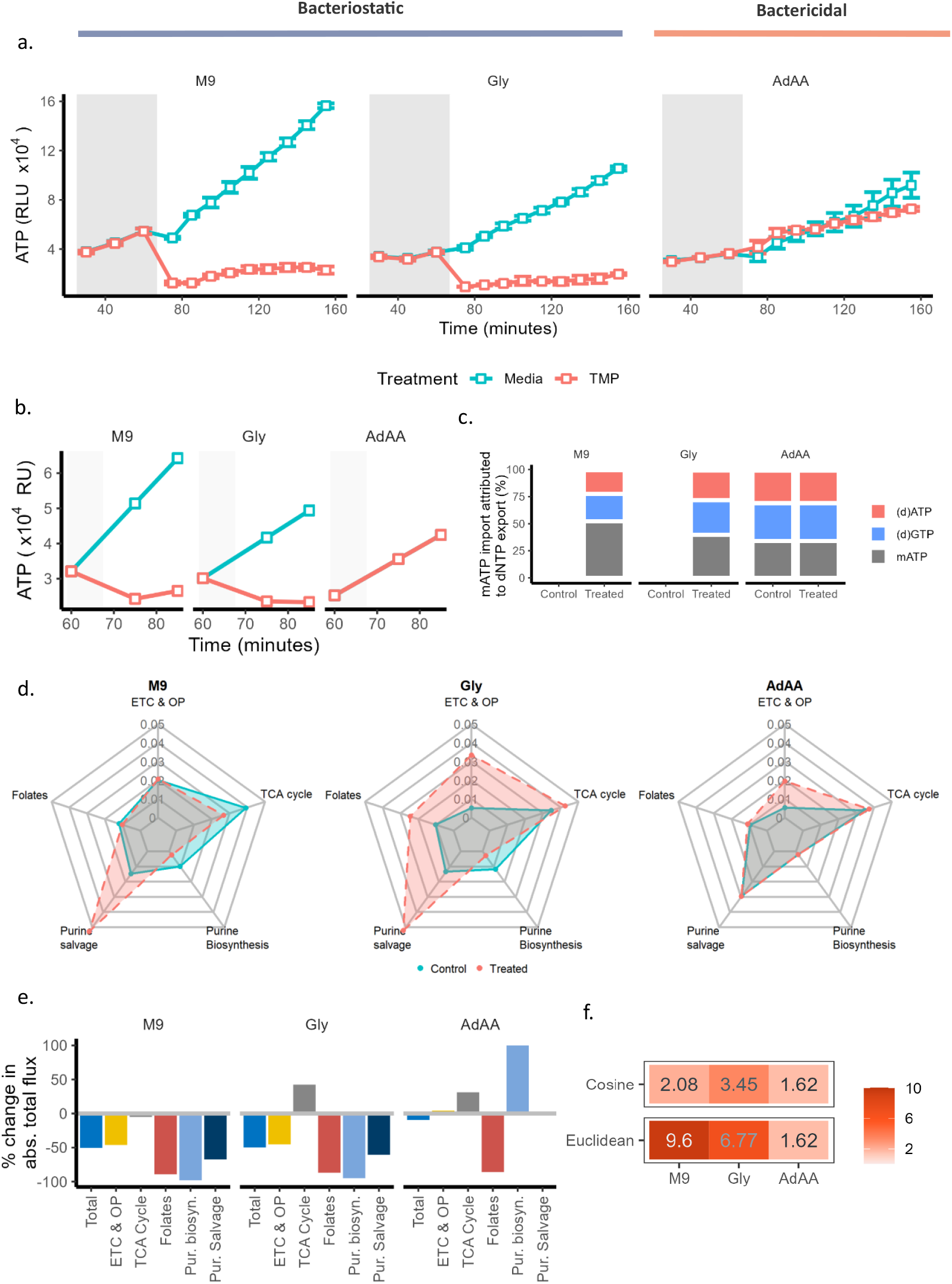
Simulated time-course ATP bioluminescence assay comparing treatment groups in different media conditions after trimethoprim-mimicking stress at the 75-minute mark. (A) Time-course ATP bioluminescence assay of a trimethoprim (TMP) susceptible clinical UPEC isolate. Assay was performed in triplicates in M9 minimal media which was either not supplemented (M9), supplemented with glycine (Gly), or with adenine, glycine, and methionine (AdAA). The grey box denotes baseline measurements before treatment of blank media or trimethoprim was added. Each data point is the mean of 4 biological replicates with error bars showing standard error mean of 95% confidence intervals. (B) Simulated change in total free ATP (relative units; RU) over time. Grey box indicates baseline measurements with weighting applied to folate associated reactions at 75 minutes for treated groups (TMP; trimethoprim). (C) Proportion of mATP import flux attributed to the export of a given nucleotide. (D) Centrality (PageRank) of relevant pathways. Values given at 75 minutes per media group on a radar chart where each ring is 0.01 step and coloured by treatment group. Solid blue lines and shading denote control group and dashed red line with shading denote trimethoprim-treated group. (E) Percentage change in absolute total flux between treated and control groups at 75 minutes in relevant pathways. (F) Cosine and Euclidean distance between flux vectors from LP solutions at 75 minutes between control and treated groups. Cosine distance was multiplied by 100 and Euclidean distance divided by 10,000 for clarity.

The switch from *de novo* purine biosynthesis to salvaging mATP resulted in significant network-wide changes in bacteriostatic conditions. M9 media conditions had the smallest change in reaction sets (suggested by cosine distance) compared to the other media compositions yet had the greatest change in flux within the network (suggested by Euclidean distance) (Figure 4f). Whereas the change in reaction sets and overall flux between control and treated groups in M9+Gly and AdAA conditions were comparable and appear to be attributed to the shift from glycine to glucose and ammonium.

When constraints were applied to the folate reactions, this coincided with a decline in the absolute total flux through folate associated reactions (Table S5) in all media conditions (M9: -91.0%, M9+Gly: -63.6%, M9+AdAA: -68.1%; Figure 4e).

### Role of glycine in trimethoprim challenge

The role of glycine in trimethoprim challenge has remained elusive; we sought to understand its function by clustering pathways on the flow of mass through the network, thereby separating reactions into functional groups. With this approach, we identified glycine as the one-carbon unit donor for THF, amino acids, and pyruvate during trimethoprim challenge (Figure 5). In untreated conditions lacking glycine (bacteriostatic), the metabolites pyruvate and ammonium were used to synthesise serine which acted as the one-carbon unit donor for THF which also produced glycine via serine hydroxymethyltransferase. In this scenario, glycine was not recycled by the glycine cleavage system to provide a second one-carbon unit to THF. Instead, it was used as a carbon source to make purines, threonine, and other amino acids that were synthesised from glutamate such as isoleucine (Figure 5). When folate availability was restricted by applying weightings to folate associated reactions, the solution had reduced glycine usage; glycine was no longer involved in threonine, amino acid, and purine biosynthesis. Any glycine that was needed (such as biomass glycine for proteins) was produced via serine-glyoxylate aminotransferase.

**Figure 5.**
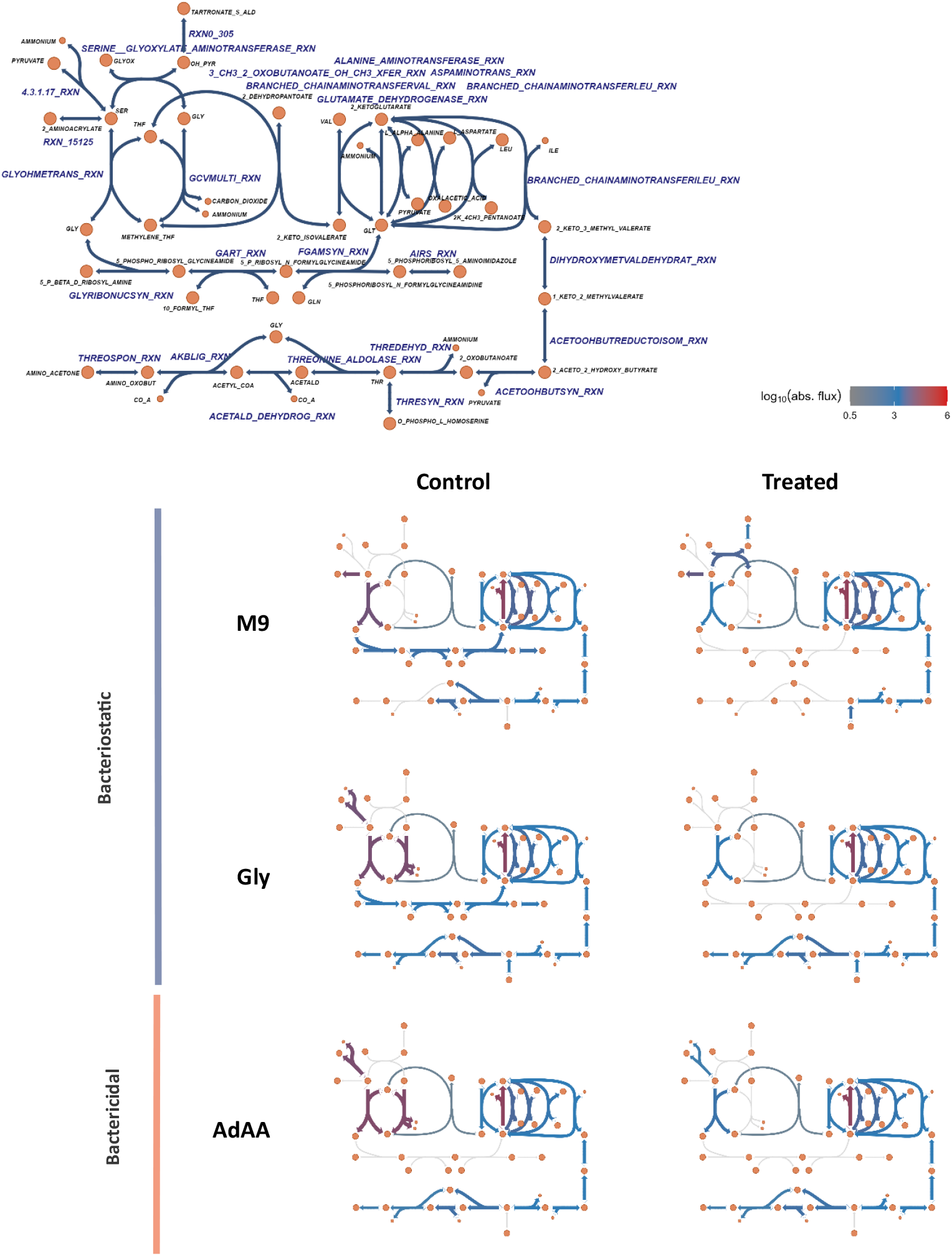
Map of glycine utilisation during control and trimethoprim-mimicking conditions at the 75-minute mark under different media conditions. (top) Network map of reactions from communities that contained a reaction involving glycine which includes metabolite and reaction names. (bottom) Unlabelled network maps with colour of reaction proportional to the log^10^ absolute flux of that reaction in each scenario.

In all scenarios where glycine was available in the media, the solution involved utilising glycine to synthesise pyruvate following the L-threonine degradation III pathway and super-pathway of methylglyoxal degradation (Figure 5). When untreated, serine was still produced from pyruvate and ammonium for serine hydroxymethyltransferase to donate one-carbon units to THF. However, the glycine cleavage system was active, and glycine was directly used to synthesise threonine, other amino acid synthesis, and pyruvate via glycine C-acetyltransferase and threonine aldolase. This suggests glycine was an effective precursor for these compounds but too expensive to synthesise *de novo* as a precursor. Trimethoprim-mimicking conditions did little to influence the utilisation of glycine; this removed the utilisation of the glycine cleavage system and reduced serine hydroxymethyltransferase. All other uses of glycine were preserved. The addition of adenine and methionine (bactericidal) had no effect on the utilisation of glycine except that glycine was not used as a carbon donor for purine biosynthesis due to adenine salvage (Figure 5).

### Experimental validation of ATP salvage during trimethoprim challenge

Experimental validation of modelling results was performed by screening ninety-six randomly selected clinical UPEC isolates (equal numbers of trimethoprim susceptible and resistant) to determine whether trimethoprim leads to energy exhaustion in bacteriostatic conditions. In M9 minimal media (bacteriostatic), there was an average of 29.7% reduction in intracellular ATP for trimethoprim susceptible UPEC compared to a 3.4% increase in intracellular ATP for resistant isolates (p<0.001, CI [0.10, 0.07]) (Figure 6a&b). This suggests that susceptible populations are salvaging ATP in response to trimethoprim exposure to a degree that can distinguish them from resistant populations in bacteriostatic conditions (ROC-AUC: 0.817) (Figure 6c). There were two optimal cut-off values of -30% (TPR 0.625, TNR 0.937) and -11% (TPR 0.812, TNR 0.75) for distinguishing these populations, the latter of which had 78.1% agreement with the phenotypic results provided by the Clinical Microbiology Laboratory that provided the isolates. Further, of the media compositions tested, M9 (bacteriostatic) provided the most consistency between repeated sampling with 2σ in susceptible and resistant groups being ±66.3% and ±51.9%, respectively (Figure S5). The supplementation of glycine was comparable to base M9 media with a mean decline of 28.6% in intracellular ATP in susceptible isolates compared to a 1.8% increase for resistant isolates (p<0.001, CI [0.12, 0.06]) (Figure 6a&b). Likewise, accuracy in distinguishing phenotypic populations was comparable (ROC-AUC 0.81) at its optimal cut-off of -15% (TPR 0.79, TNR 0.87). In contrast, the presence of adenine, glycine, and methionine which elicited a bactericidal outcome did not lead to significant ATP decline in both phenotypic populations (ROC-AUC 0.36). There was a 14.3% mean increase in ATP in the susceptible group compared to a 4.5% decline for resistant isolates (p = 0.01, CI [0.12, 0.05]).

**Figure 6.**
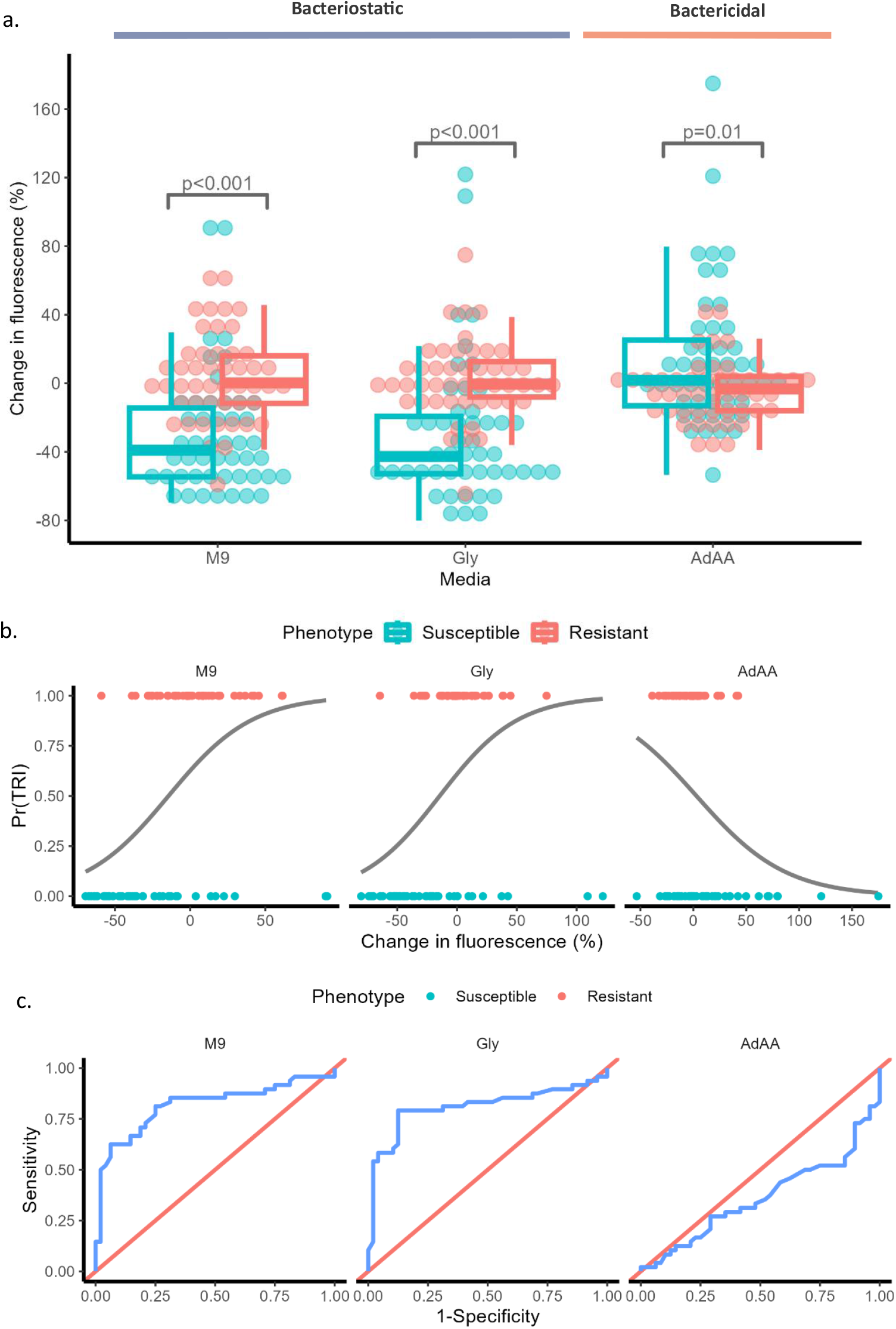
Measuring change in bacterial ATP following 30 minutes of trimethoprim exposure in based M9 media and with supplementations of glycine (Gly) or adenine, glycine, and methionine (AdAA) comparing trimethoprim susceptible and resistant populations. (A) boxplot illustrating the mean percentage change (n=4) in intracellular ATP between an untreated control and a treated sample for 96 clinical UPEC isolates (48:48 susceptible [S] to resistant [R]) after 30 minutes of exposure to trimethoprim. This was performed in triplicates for the same media conditions as the time-course assay. (B) Binomial logistic regression curve for probabilities of trimethoprim resistance, Pr(TRI), against percentage change in fluorescence between a treated and untreated sample by media condition. Grey line is predicted values. (C) Receiver operating characteristic plot for (B) illustrating the true positive rate (TPR or sensitivity) and false positive rate (FPR) subtracted from 1 (specificity) over a range of cut-off values for differentiating trimethoprim susceptibility. The diagonal line represents the chance diagonal.

## Capacity of different trimethoprim resistance determinants in limiting ATP salvage

As resistance to trimethoprim was demonstrated to provide protection against energy exhaustion in bacteriostatic conditions (Figure 6), we sought to investigate whether different resistance determinants provided the same degree of protection (Figure S6). Given bactericidal conditions did not lead to energy exhaustion in susceptible groups, we focused on the two bacteriostatic conditions – M9 with and without glycine supplementation. A total of 42 isolates with known *dfrA* variants were investigated which carried either: *dfrA1* (n=16, M9=6.5%±27.9, Gly=-0.28%±18.7), *dfrA17* (n=12, M9=1.3%±18.4, Gly=5.8%±18.8), *dfrA14* (n=5, M9=1.3%±17.6, Gly=7.9%±15.5), *dfrA12* (n=4, M9=22.9%±33.2, Gly=16.1%±41.6), *dfrA5* (n=3, M9=-5.2%±28.7, Gly=-5.9%±20.2), or *dfrA1*+*dfrA12* (n=2, M9=-0.58%±12.9, Gly=-14.4%±20.8) (Figure S6). Occurrences of only a single *dfrA* type were excluded: *dfrA15* (M9=33.1%, Gly=-8.4%), *dfrA36* (M9=29.4%, Gly=12.5%), and *dfrA1*+*dfrA14* (M9=-1.7%, Gly=23%) (Figure S6). There were three isolates with no known trimethoprim resistance mechanism and no trimethoprim susceptible isolate carried a known resistance mechanism. The mean change in fluorescence in both media for 5/6 *dfrA* variants tested were within ±10% with exception of *dfrA12* which skewed towards a larger positive value.

## Discussion

An *E. coli* UTI89 GSM model of trimethoprim-induced constraint on purine biosynthesis supported the hypothesis of free ATP being used to restore the purine and nucleoside availability in bacteriostatic conditions. This was due to trimethoprim disrupting the purine biosynthesis pathway which is a major source of *de novo* ATP. In bactericidal conditions, the availability of a purine in the media negated this effect and free ATP remained untouched. These findings match the observations from *in vivo* metabolomics and with other antibiotics ^1,31,32^. The possibility of ATP being used for functions not considered in the model (e.g., motility, stress response, and repair) cannot be excluded. The synthesis of metabolites identified here as inhibited by trimethoprim agree with the metabolomics observations of Kwon *et al*. ^1^. However, the approach used here could not determine if metabolites accumulated or depleted, so only metabolites that directly involved folates for biosynthesis were identified. For example, our results suggested trimethoprim inhibits the synthesis of co-factors NAD and FAD in bacteriostatic conditions, but there are no known reports of this *in vivo*.

It is possible that any effects on the availability of NAD and FAD requires multiple generations, by which time the stalling of DNA, RNA, and protein biosynthesis prohibits cellular replication. However, this may not be the case for sulfonamides given bacteria require multiple generations before sulfonamides have an effect ^9,33^.

The model supported that the availability of methionine restored its restricted synthesis and restored spermidine. However, the model does not exclude the likelihood of methionine’s role as the protein synthesis initiator, in the form of formyl-methionyl-tRNA^met^, being relevant to bactericidal activity. To better understand the flow of glycine through the network, a combination of mass flow graphs and the Leiden algorithm were used to divide the network into functional communities. This further divided the network into sets of reactions where there were notable changes between treatment groups which enabled identification of where glycine was being used. We found that when glycine is available in the environment, the network used it to synthesise pyruvate which could serve to produce ATP and offer protection against reactive oxygen species ^34^.

The cause of the depletion in intracellular ATP for trimethoprim susceptible *E. coli* in M9 minimal media challenged with trimethoprim could arguably be attributed to the bacteriostatic behaviour of trimethoprim in this condition. With the arrest of DNA, RNA, and protein synthesis caused by trimethoprim, and without metabolites available to compensate for the effect, the demand for ATP would decline. This agrees with the observations in *E. coli* by Lobritz et al. ^35^ whereby bacteriostatic antibiotics slow down cellular respiration within 10 minutes of exposure. Nonetheless, the drop in intracellular ATP observed here and by Kwon *et al*. ^1^ suggest cellular arrest of catabolic processes which in turn slows down oxidative phosphorylation. This hypothesis is contradicted by an observation from Stepanek et al. ^36^ who demonstrated *Bacillus subtilis* grown on Belitzky minimal medium supplemented with at least 10 mM ATP can tolerate 20-fold higher concentrations of trimethoprim compared to without ATP supplementation. Likewise, accumulations of adenine-based metabolites like ATP at sites of infection reduce the lethal effects of antibiotics ^37^. These observations suggest bacteriostatic trimethoprim activity leads to an ATP expensive scenario rather than just cellular arrest. There is a growing body of evidence suggesting that purines, including phosphorylated purines like ATP, are depleted by trimethoprim’s bacteriostatic activity and that respiration is not the cause of the observed effect ^1,36,38^. A recent genome-scale metabolic modelling study by Yang *et al*. ^31^ explored how antibiotics affect cellular energy with experimental validation. They provided evidence to suggest that purine limitation caused by trimethoprim challenge, similarly with mutations in purine biosynthesis genes, increased ATP demand to restore the nucleoside pool.

Indeed, models of this behaviour suggested adenine supplementation dramatically reduced that ATP demand. This agrees with the observation by Kwon *et al*. ^1^ that purine supplementation did not lead to nucleoside depletion from trimethoprim challenge. The hypothesis of using ATP to restore nucleoside imbalance agrees with the observations previously mentioned, and importantly, does not contradict the observations of arrested DNA, RNA, and protein synthesis. Therefore, it would be logical to assume that the means of improving the assay’s performance is by ensuring there is a demand for nucleosides. This would explain why no depletion of intracellular ATP is observed in bactericidal trimethoprim conditions as the nucleoside pool is compensated by a purine. Previous observations suggested glycine supplementation could improve ATP decline during trimethoprim challenge, but we observed no improvement over base M9 media. This suggests that the results reported by Kwon *et al*. ^1^ in *E. coli* K-12 NCM3722 are not representative of *E. coli* and that the conclusion of glycine being the first metabolite to deplete after trimethoprim exposure is perhaps not the case for most *E. coli*.

In conclusion, the results presented here provide strong evidence that bacteriostatic trimethoprim activity disrupts purine biosynthesis with free ATP being salvaged to alleviate trimethoprim’s actions.

## Resource availability

Requests for further information and resources should be directed to and will be fulfilled by the lead contact, Cailean Carter. GSM model is available on BioModels under accession MODEL2507240001. Code is available on Zenodo (https://doi.org/10.5281/zenodo.16412170).

## Acknowledgements

We thank Mark Poolman, Sisse Mortensen, and John Elmerdahl Olsen for allowing us to use their *E. coli* UTI89 model and for training on ScrumPy. Thank you to James Lazenby for training and assistance with the CLARIOstar Plus.

CC is supported by the UKRI Medical Research Council Doctoral Antimicrobial Research Training (DART) Industrial CASE Programme [grant number MR/R015937/1], as a CASE award in collaboration with Test&Treat. GCL, JW, and DS are supported by the Biotechnology and Biological Sciences Research Council (BBSRC) Institute Strategic Programme Microbes in the Food Chain BB/R012504/1 and its constituent project BBS/E/F/000PR10349 until April 2023 before being funded under BBSRC Institute Strategic Programme Microbes and Food Safety BB/X011011/1 and its constituent project(s) BBS/E/F/000PR13634 (Theme 1, Microbial threats from foods in established and evolving food systems).

## Supplementary information

Document S1. Figures S1-S6, Tables S1-S6, and supplementary references.

## Transparency declaration

Authors declare no conflicts of interest.

