## Supplementary document for "Genome-scale metabolic modelling of trimethoprim’s mode of action reveals free ATP salvaged to restore purine pool"

### Supplementary material

##### Model properties and validation

The reactions contained within the model were modularised for ease of analysis, thereby aiding focus on or exclusion of groups of reactions. Within ScrumPy, a top-level module was loaded which imports files containing the intracellular reactions and transporters (summarised in Figure S1). The intracellular reactions included the polymerisation reactions and any manually corrected reactions (termed *E. coli*), the electron transport chain (ETC), and the cytosolic reactions generated during the construction of the model from the *E. coli* UTI89 BioCyc database v22.0. Transporter reactions encompassed the importation of M9 minimal media and atmospheric gases, and biomass export reactions that represented cellular components *E. coli* would synthesis when growing.

Fatty acid, peptidoglycan, and lipopolysaccharide synthesis pathways were condensed into a single reaction per pathway to reduce complexity. The reaction stoichiometry for each reaction is outlined below:

**Fatty acid synthesis**

8 acetyl-CoA + 14 NADPH + 7 ATP + 6 H^+^ + H_2_O ↔ palmitate + 14 NADP + 8 CoA + 7 ADP + 7 Pi

**Peptidoglycan synthesis**

3 L-α-alanine + PEP + NADPH + D-glutamate + 5 ATP + meso-diaminopimelate + Und-PP + 2 UDP-GlcNAc ↔ peptidoglycan + 5 ADP + 7 Pi + UMP + UDP + NADP

**Lipopolysaccharide synthesis**

2 CMP-Kdo + 4 ADP-L-glycero-D-manno-heptose + 3 UDP-glucose + 2 UDP-GlcNAc + 6 palmitate + 3 ATP + UDP-galactose ↔ LPS + 7 ADP + acetate + UMP + 5 UDP + 2 CMP

| **Table S1. Biomass components of *E. coli* UTI89 as described in the model. All values are in mmol/gDW with default growth rate (μ) of 1.** | | | | | |
| --- | --- | --- | --- | --- | --- |
| **Amino acids** | | **RNA** | | **Cell wall** | |
| Alanine | 0.60664 | ATP | 0.109351 | Cardiolipin | 0.003198 |
| Arginine | 0.177753 | CTP | 0.087659 | Lipopolysaccharide | 0.008367 |
| Asparagine | 0.191759 | GTP | 0.130251 | Peptidoglycan | 0.024752 |
| Aspartate | 0.191792 | UTP | 0.094478 | Phosphatidate | 0.032487 |
| Cysteine | 0.078999 | **DNA** | | Phosphatidylethanolamine | 0.029767 |
| Glutamine | 0.188763 | dATP | 0.016927 | Phosphatidylglycerol | 0.028573 |
| Glutamate | 0.188794 | dCTP | 0.017002 | Phosphatidylserine | 0.0281 |
| Glycine | 0.862496 | dGTP | 0.015649 | **Other** | |
| Histidine | 0.065315 | dTTP | 0.015949 | Putrescine | 0.227814 |
| Isoleucine | 0.231768 | **Cofactors** | | Spermidine | 0.028084 |
| Leucine | 0.360529 | Acetyl-CoA | 0.000214 | Starch | 0.138769 |
| Lysine | 0.244788 | CoA | 0.000135 |  |  |
| Methionine | 0.109425 | FAD | 0.000176 |  |  |
| Phenylalanine | 0.119286 | 5-methyl THF | 0.000904 |  |  |
| Proline | 0.200492 | NAD | 0.001665 |  |  |
| Serine | 0.214286 | NADH | 0.000042 |  |  |
| Threonine | 0.222253 | NADP | 0.000093 |  |  |
| Tryptophan | 0.030324 | NADPH | 0.000279 |  |  |
| Tyrosine | 0.080788 | Succinyl-CoA | 0.00007 |  |  |
| Valine | 0.124954 |  |  |  |  |
| Abbreviation: gDW, gram dry weight. | | | | | |

Transporter reactions were added for the import of adenine, glycine, methionine, and thymidine from the media. For handling mATP, four reactions were added. One reaction converted ATP to mATP at no cost and one reaction transported mATP outside the system. The other two reactions were the same as described but in reverse to track import and export of mATP (Table S2). These reactions were only included in the analysis when specified.

| **Table S2.** **Summary of reactions involved with mATP** | |
| --- | --- |
| **Reaction name** | **Stoichiometry** |
| mATP_tx (importer) | x_mATP → mATP |
| mATPSynth-RXN | ATP → mATP |
| mATPase-RXN | mATP → ATP |
| mATP_bm_tx (exporter) | x_mATP ← mATP |

Some corrections were made during the validation process. Firstly, the model incorrectly contained the reaction “AIRCARBOXY-RXN” which added a carbon unit from carbon dioxide to the purine ring. In *E. coli*, this reaction was performed by two separate enzymes and used bicarbonate as the carbon donor for the purine ring according to the *E. coli* UTI89 EcoCyc v27.1 database. Reactions “RXN0-742” and “RXN0-743” which perform this step were already present in the model, thus, removing “AIRCARBOXY-RXN” from the model was the only action taken to address this. The sulphite dehydrogenase reaction was changed from reversible to forward direction to conserve internal energy balance and match the MetaCyc description of the reaction (accessed 20/06/2023). Two reactions were added to the ETC module to match the *E. coli* UTI89 EcoCyc v27.1 database:

**1.10.2.2-RXN**

Ubiquinols + 2 Cytochromes-C-Oxidized + 2 H^+^ → 4 x_H^+^ + Ubiquinones + 2 Cytochromes-C-Reduced

**RXN0-5268**

O_2_ + 2 Ubiquinols + 8 H^+^ → 8 x_H^+^ + 2 H_2_O + 2 Ubiquinones

The metabolic model encompassed 1265 reactions for the conversion of 1216 metabolites. However, 44.6% of reactions were unable to carry flux under steady state (namely dead reactions). Similarly, 45.1% of metabolites could not be balanced under steady state (namely orphan metabolites), these could only be produced or consumed. Nonetheless, a significant portion of dead reactions and orphan metabolites was expected due to the incomplete information available for any given bacterial organism. Similarly, the automated process of building a GSM from whole genomes included the errors and inconsistencies present in these databases and needed to be manually curated. Several aspects of the model were checked for consistency and validation (Table S3):

- Atomic balance: checked the proportion of atoms (C, N, P, S, O, and H) in substrates matched that in products for each reaction.
- Energy and redox conservation: checked ATP, NADH and NADPH were not produced in the absence of flux into the network.
- Mass conservation: similarly, checked the model was not producing compounds from nothing.
- Stoichiometry inconsistencies: checked reactions were balanced. For example, if two reactions with the same reactants and products were producing different quantities of products from the same quantity of reactants.
- Electron transport chain: checked the model produced energy from compounds in the media as expected.
- Biomass production: checked the model could synthesise all necessary biomass components.

Following these steps, the model could successfully synthesise all biomass components (Figure S2).

| **Table S3. Properties of the *E. coli* UTI89 GSM model** | |
| --- | --- |
| **Quantitative properties of model** | **Total** |
| Reactions (excluding transporters) | 1265 |
| Transporters | 77 |
| Reactions associated with identifiable genes | 1006 |
| Metabolites | 1216 |
| Dead reactions | 564 |
| Orphan metabolites | 548 |
| Unbalanced reactions | 0 |
| Metabolites with undefined empirical formula | 0 |
| **Qualitative properties of model** | **Checked?** |
| Atomic balance for C, N, S, P, O, H | Yes |
| Energy conservation | Yes |
| Biomass production | Yes |

| **Table S4. Reactions grouped into their respective pathways.** Reactions are their common name or BioCyc ID when common name not available. | |
| --- | --- |
| **Pathway** | **Reactions** |
| Folates | Table S5 |
| Purine biosynthesis | Amidophosphoribosyltransferase, phosphoribosylglycinamide formyltransferase, phosphoribosylformylglycinamide cyclo-ligase, phosphoribosylaminoimidazole-succinocarboxamide synthase, phosphoribosylamine—glycine ligase, AICAR transformylase, IMP cyclohydrolase, adenylosuccinate lyase, adenylosuccinate lyase, N^5^-carboxyaminoimidazole ribonucleotide mutase, 5-(carboxyamino)imidazole ribonucleotide synthase, phosphoribosylglycinamide formyltransferase 2, phosphoribosylformylglycinamide synthetase |
| Purine salvage | Adenine deaminase, adenosine deaminase, nucleoside phosphorylase, deoxynucleoside kinase, purine nucleoside phosphorylase, adenylate kinase, nucleoside diphosphate kinase, ribonucleoside-diphosphate reductase 1, guanylate kinase, GMP reductase, guanine deaminase, guanylate kinase, inosine/guanosine kinase, xanthine dehydrogenase, adenylosuccinate lyase, deoxyadenosine kinase, AMP phosphorylase, ribonucleoside hydrolase, adenylosuccinate synthetase, adenine phosphoribosyltransferase, guanine phosphoribosyltransferase, nucleoside triphosphate pyrophosphohydrolase, dGTP triphosphohydrolase, deoxyadenylate kinase |
| Tricarboxylic acid cycle | Branched-chain α-keto acid dehydrogenase complex, lipoamide dehydrogenase, branched-chain α-keto acid dehydrogenase complex, 2-oxoglutarate decarboxylase, 2-oxoglutarate synthase, aconitate hydratase A, citrate synthase, fumarase, isocitrate dehydrogenase, malate thiokinase, malate dehydrogenase, aconitate hydratase A, succinate dehydrogenase, succinate:caldariellaquinone oxidoreductase complex, dihydrolipoyltranssuccinylase, succinyl-CoA synthetase, succinate:quinone oxidoreductase, malyl-CoA lyase, malate-CoA ligase, isocitrate lyase, aspartate aminotransferase, glutamate dehydrogenase, glutamate synthase, 2-methylcitrate synthase, 2-methylcitrate dehydratase, 2-methylisocitrate lyase, fumarate reductase, succinyl-diaminopimelate desuccinylase, phosphoserine/phosphohydroxythreonine aminotransferase, tetrahydrodipicolinate succinylase, |
| Electron transport chain & oxidative phosphorylation | NADH:quinone oxidoreductase, NADH:menaquinone oxidoreductase, cytochrome-c-oxidase, succinate dehydrogenase, cytochrome-c-reductase, cytochrome-b-oxidase, ATP synthase, nitrate reductase (quinone), alcohol dehydrogenase (quinone) |

| Table S5. List of folate associated reactions. Reaction names are their common names when available, otherwise a BioCyc ID is used. | |
| --- | --- |
| Reaction name | **Stoichiometry** |
| AICAR transformylase | AICAR + 10-formyl-THF ↔ FAICAR + THF |
| Methylene THF dehydrogenase | NADP + methylene-THF ↔ NADPH + 5,10-methenyl-THF |
| Dehydropantoate hydroxymethyltransferase | H_2_O + 2-keto-isovalerate + methylene-THF ↔ 2-dehydropantoate + THF |
| Phosphoribosylglycinamide formyltransferase | GAR + 10-formyl-THF ↔ H^+^ + FGAR + THF |
| RXN-2881 | Formaldehyde + THF ↔ H_2_O + methylene-THF |
| Methionine synthase | L-homocysteine + 5-methyl-THF → L-methionine + THF |
| 1.5.1.20-RXN | NAD + 5-methyl-THF ← H^+^ + NADH + methylene-THF |
| Glycine cleavage system | NAD + THF + glycine → CO_2_ + NH_4_^+^ + NADH + methylene-THF |
| Dihydrofolate reductase | THF + NADP ← H^+^ + NADPH + DHF |
| 1.5.1.15-RXN | NAD + methylene-THF ↔ 5,10-methenyl-THF + NADH |
| Thymidylate synthase | dUMP + methylene-THF → DHF + dTMP |
| Serine hydroxymethyltransferase | L-serine + THF ↔ H_2_O + glycine + methylene-THF |
| Methylene THF cyclohydrolase | 5,10-methenyl-THF + H_2_O ↔ H^+^ + 10-formyl-THF |
| Ferredoxin-dependent methylene-THF reductase | 5-methyl-THF + 2 Oxidized-ferredoxins ← 2 H^+^ + 2 Reduced-ferredoxins + methylene-THF |
| RXN0-1862 | UDP-4-amino-4-deoxy-L-arabinopyranose + 10-formyl-THF ↔ H^+^ + UDP-4-deoxy-4-formamido-β-L-arabinopyranose + THF |
| RXN-13909 | NAD + 10-formyl-THF → H^+^ + 10-formyl-7,8-DHF + NADH |
| 5-formyl-THF cyclo-ligase | 5-formyl-THF + ATP → 5,10-methenyl-THF + Pi + ADP |
| Trimethylsulfonium methyltransferase | Trimethyl sulfonium + THF ← H^+^ + 5-methyl-THF + dimethyl sulphide |
| Formate—THF ligase | Formate + ATP + THF → Pi + 10-formyl-THF + ADP |
| DHF synthetase | L-glutamate + ATP + 7,8-dihydropteroate → H^+^ + DHF + Pi + ADP |
| Abbreviations: dUMP, deoxyuridine monophosphate; Pi, inorganic phosphate | |

| **Table S6. Luciferase ATP bioluminescence reaction stoichiometry ^1^.** | |
| --- | --- |
| luciferase + ATP + D-luciferin → luciferase(luciferyl-adenylate) + PPi | (1) |
| luciferase(luciferyl-adenylate) + O_2_ → luciferase(oxyluciferin; AMP) + CO_2_ | (2) |
| luciferase(oxyluciferin; AMP) → luciferase(oxyluciferin; AMP) + photon | (3) |
| luciferase(oxyluciferin; AMP) → luciferase + oxyluciferin + AMP | (4) |
| ATP + O_2_ + D-luciferin → oxyluciferin + AMP + PPi + CO_2_ + photon | (Net) |

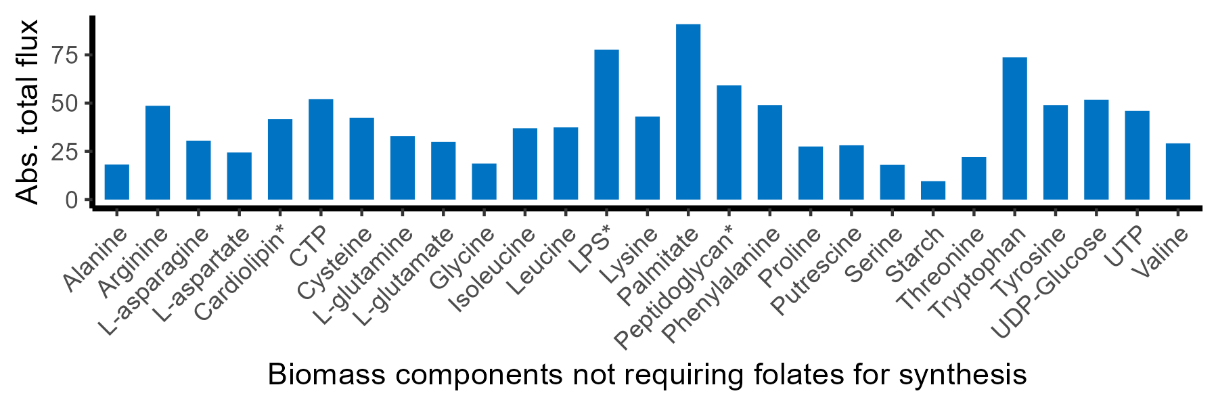

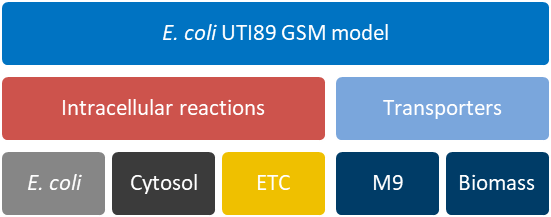

**Figure S1. Modular structure of *E. coli* UTI89 GSM model.**

**Figure S2. Biomass components not affected by trimethoprim-induced disruption of the folate pool.** Bar chart of absolute total flux of LP solution to synthesise 1 unit of biomass component. Thus, asserting the model can synthesise all biomass components.

*The absolute total flux for these metabolites was divided by 10 to improve visibility of the other metabolites on the same scale.

E. coli UTI89 GSM model

Intracellular reactions

E. coli

Cytosol

ETC

Transporters

M9

Biomass

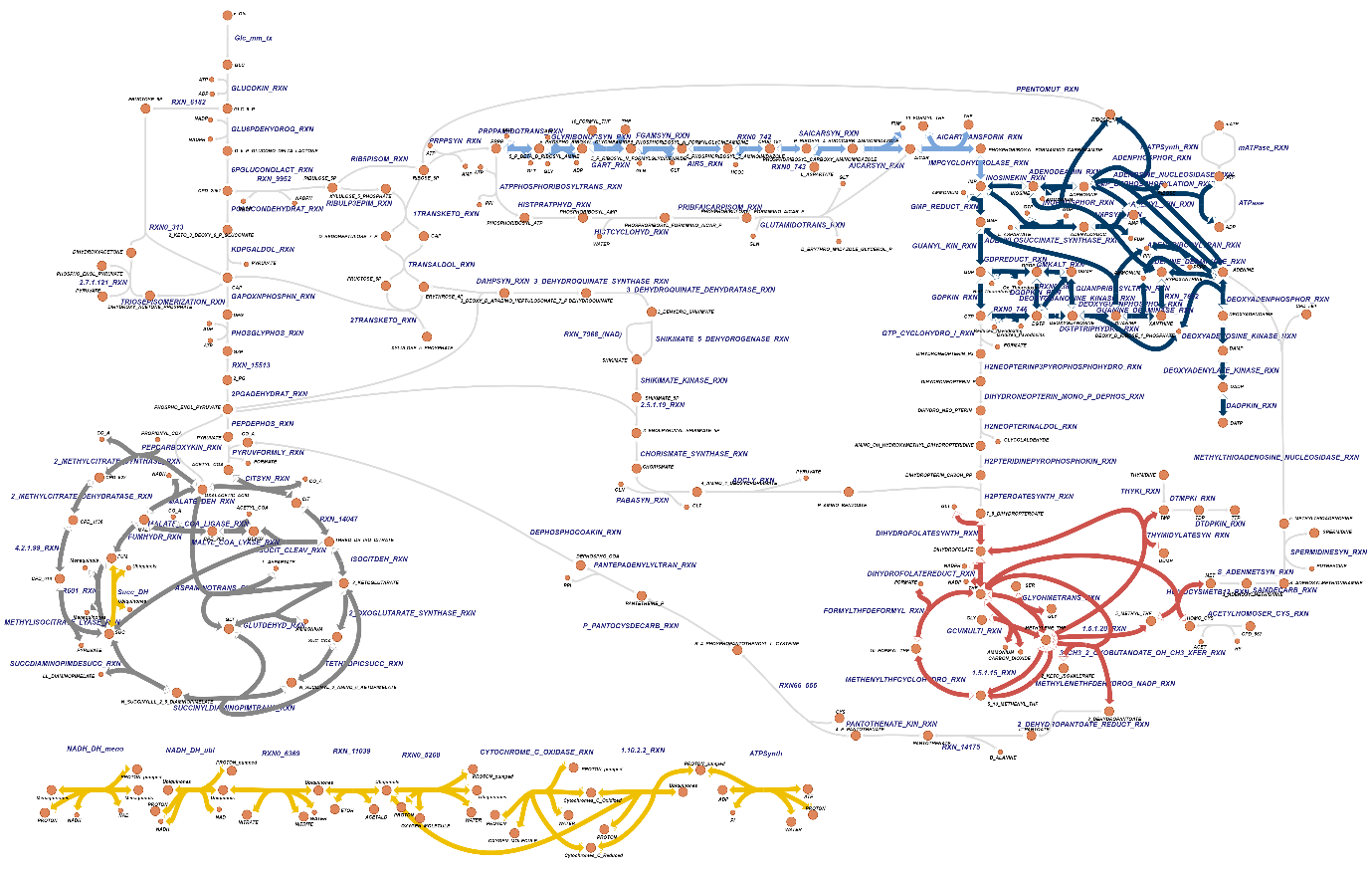

**Figure S3. Network map of core metabolism in *E. coli* UTI89 GSM model.** Electron transport chain (ETC) and oxidative phosphorylation (OP), tricarboxylic acid (TCA) cycle, folate associated reactions, purine biosynthesis, and purine salvage highlighted.

**Figure S4. Preliminary incorporation of omics data.** (A; top) Time-course absorbance assay of a trimethoprim (TMP) susceptible clinical UPEC isolate. Assay was performed in triplicates in M9 minimal media which was either not supplemented (M9), supplemented with glycine (Gly), or with adenine, glycine, and methionine (AdAA). The grey box denotes baseline measurements before treatment of blank media or trimethoprim was added. Each data point is the mean of 4 biological replicates with error bars showing standard error mean of 95% confidence intervals. (A; mid) Simulated production of biomass ATP over time. (A; Bottom) Production of total ATP over time as determined by the rate of ATP synthesis from ATP bioluminescence. In both figures, the growth rate in control conditions was used for baseline time points (≤60 mins) for the treated groups. (B) Heatmap of proportion of total ATP that is maintenance ATP (mATP) from A bottom plot. (C) Change in mATP import flux as increased weighting was applied to the mATP importer. Change in flux for mATP import with increased weighting applied to folate associated reactions when optimum mATP import weighting was applied.

*In vivo*

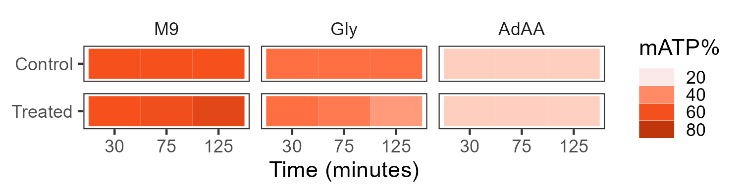

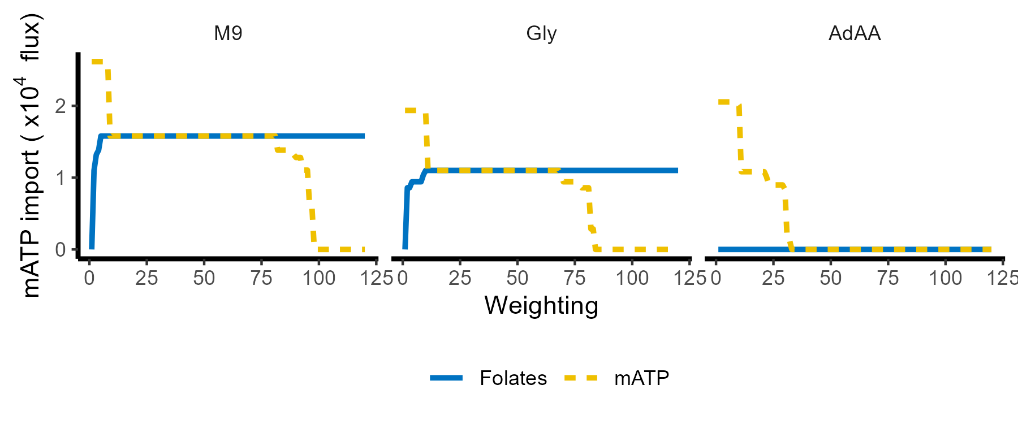

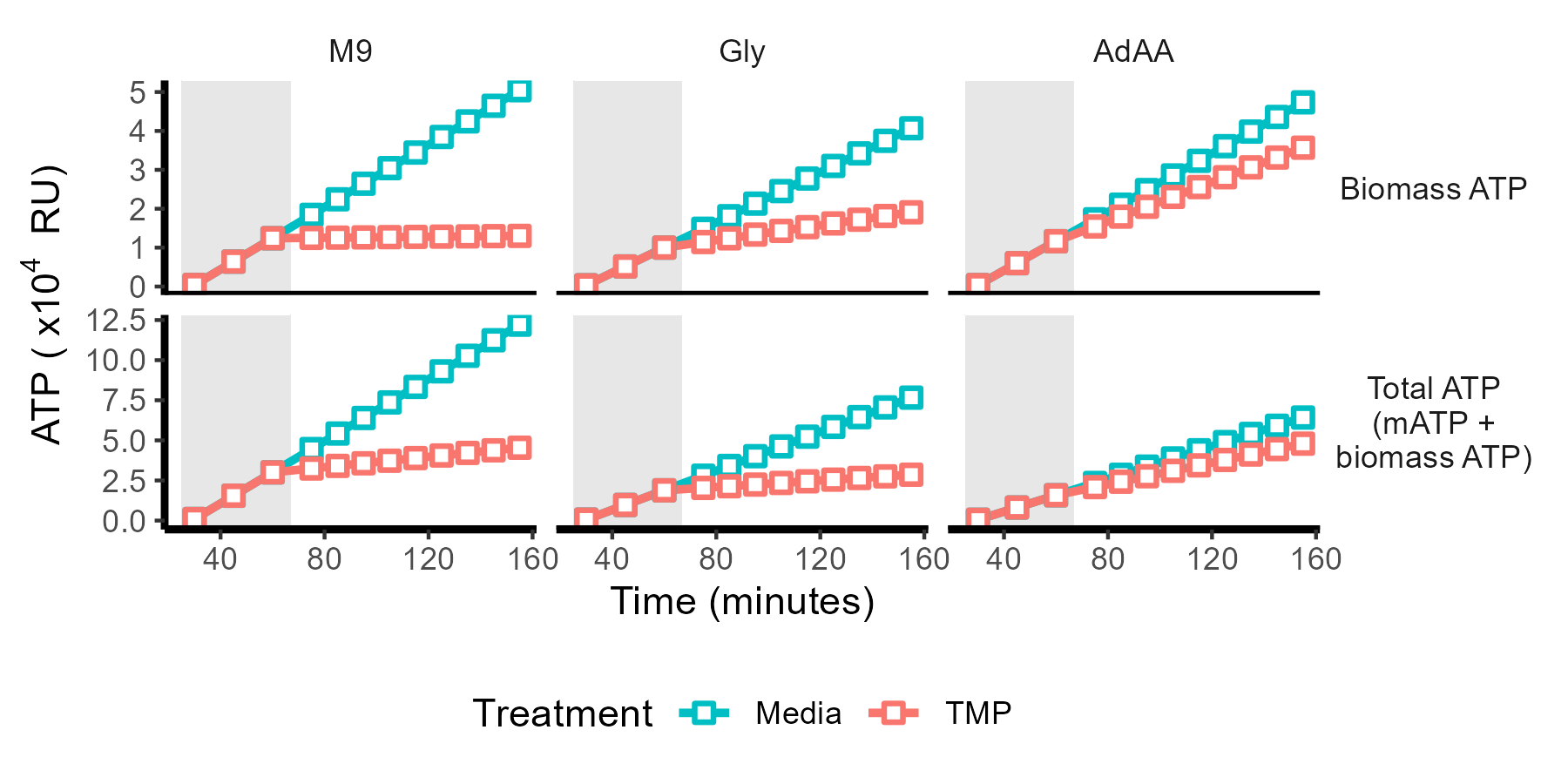

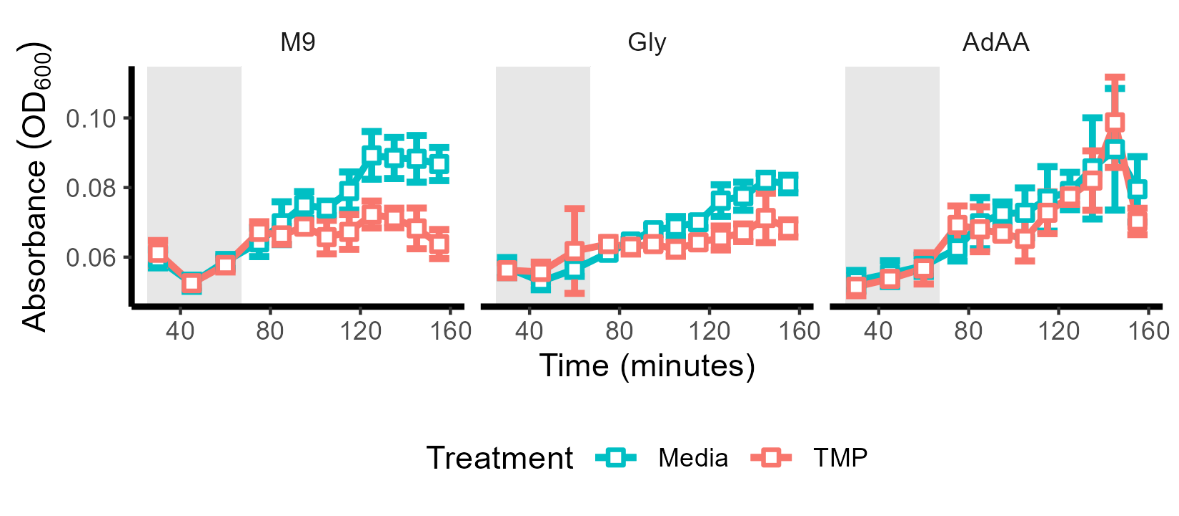

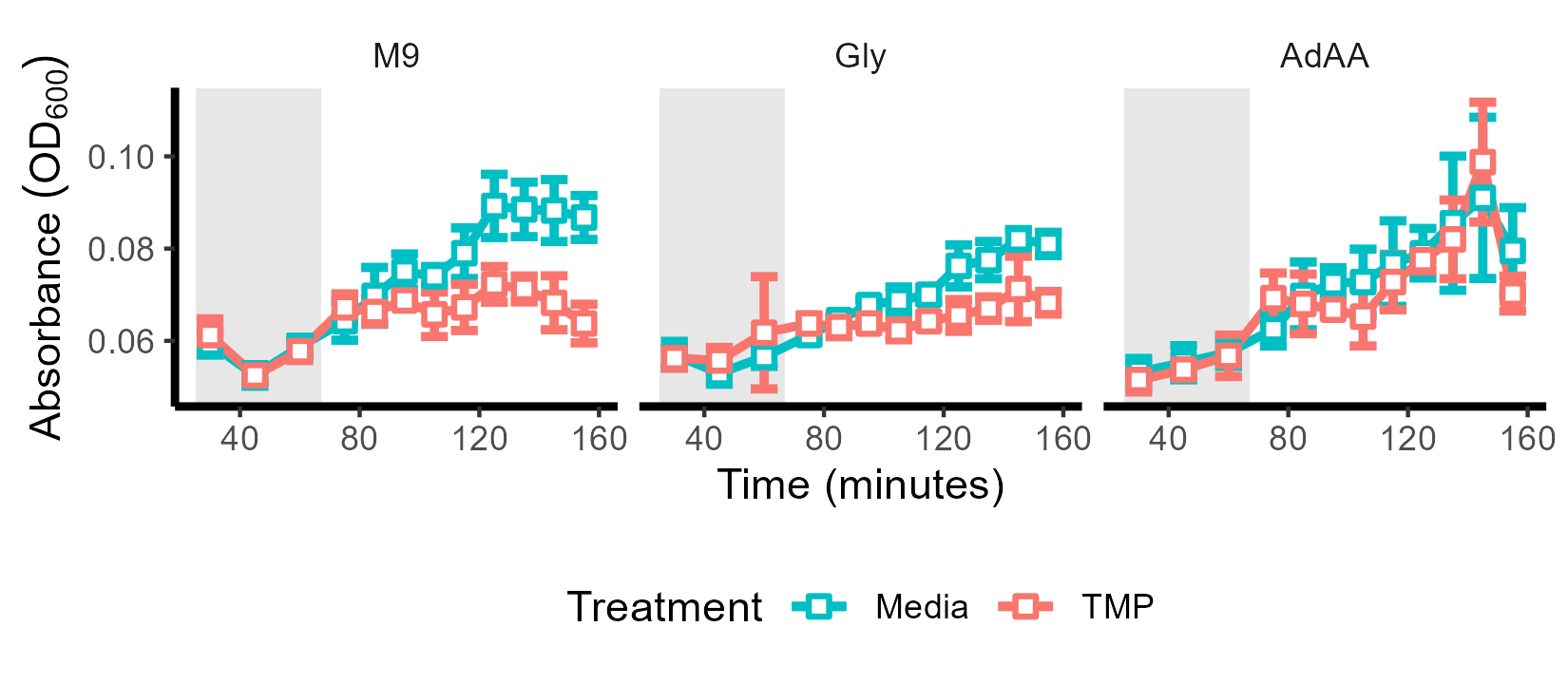

**Bacteriostatic**

**Bactericidal**

a.

b.

c.

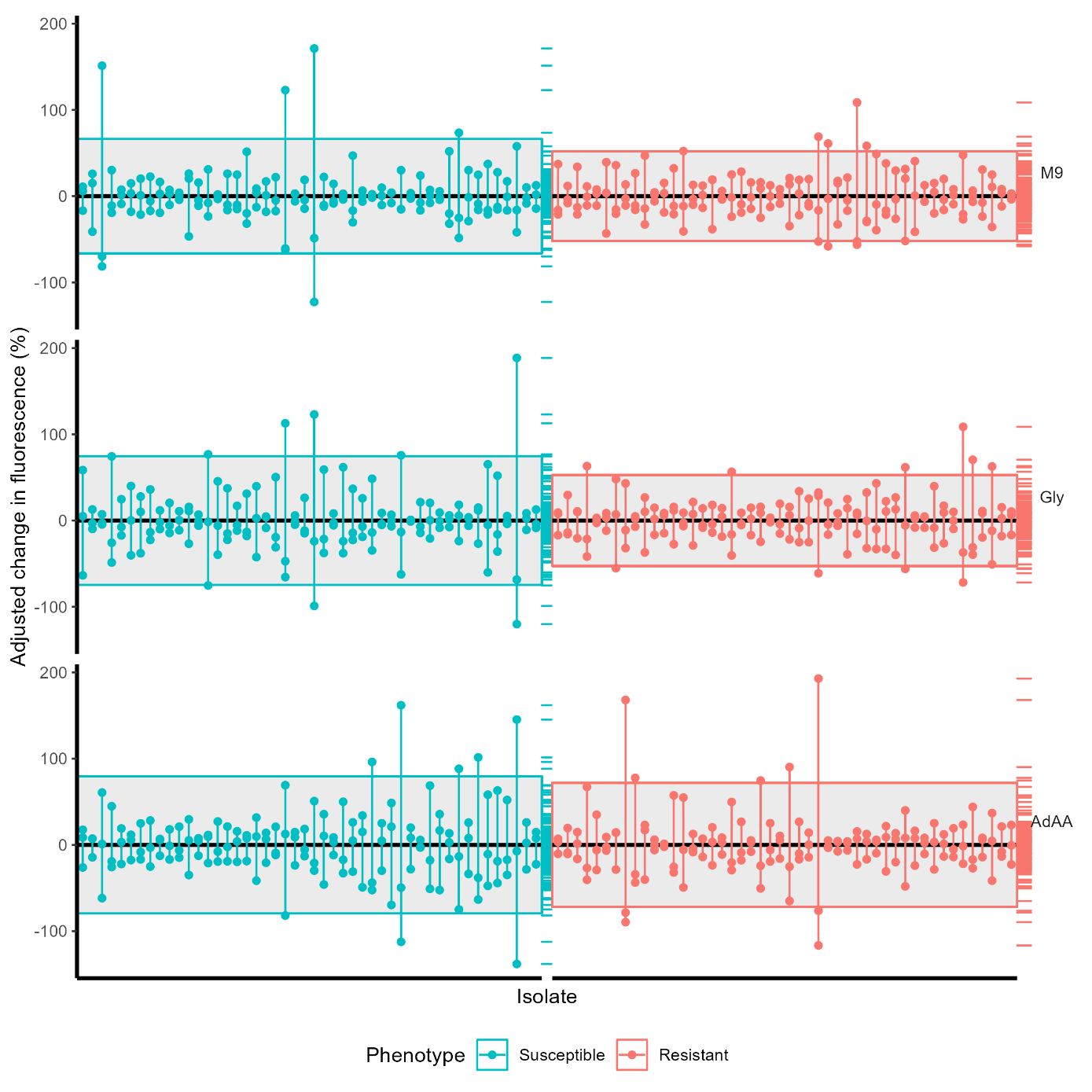

**Figure S5. Distribution in change in fluorescence of three replicates per isolate.** Each isolate is plotted along the x-axis with a vertical line between the three replicates. The black horizontal line indicates the population mean of zero. The rug plot on the right of each scatter plot illustrates data density across the y-axis. The grey box denotes 2 standard deviations ($\boldsymbol{\pm2}\boldsymbol{\sigma}$).

**Figure S6. Histogram comparing different trimethoprim resistance mechanisms in relation to change in fluorescence in bacteriostatic conditions.** NI denotes isolates where no known trimethoprim resistance determinant was identified.

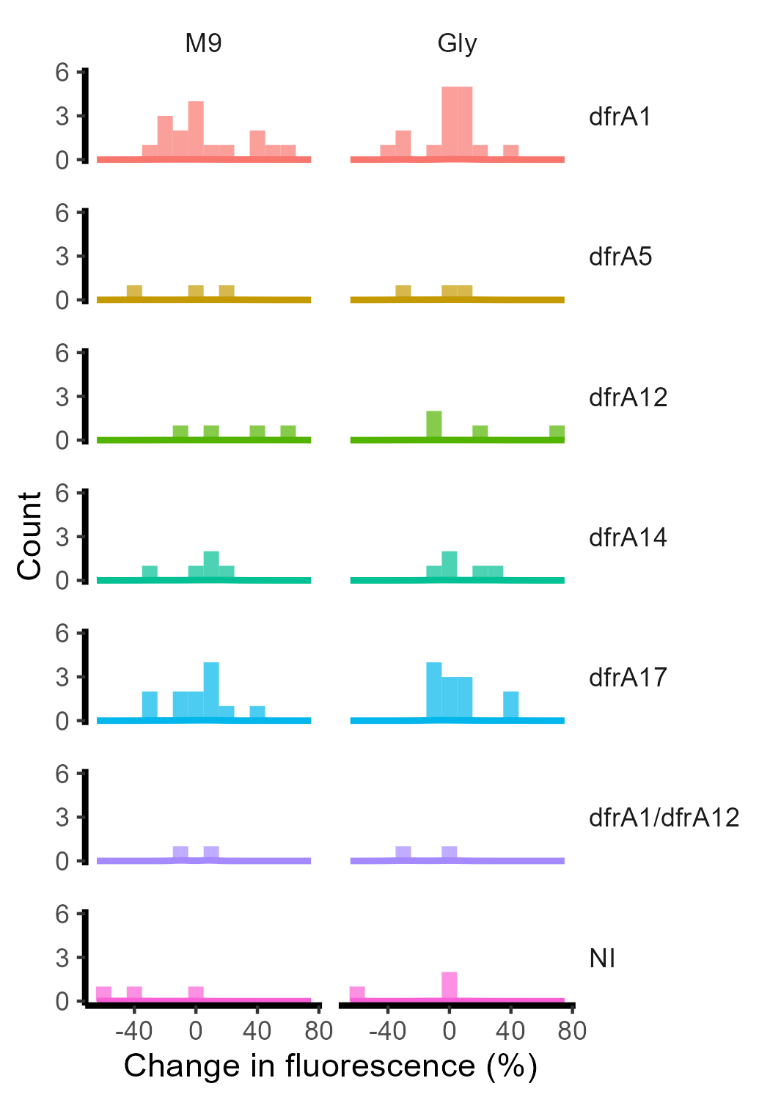
